# Interactive Visualization of Metric Distortion in Nonlinear Data Embeddings using the distortions Package

**DOI:** 10.1101/2025.08.21.671523

**Authors:** Kris Sankaran, Shuzhen Zhang, Chenab, Marina Meilă

## Abstract

Nonlinear dimensionality reduction methods like UMAP and ***t***-SNE can help to organize high-dimensional genomics data into manageable low-dimensional representations, like cell types or differentiation trajectories. Such reductions can be powerful, but inevitably introduce distortion. A growing body of work has demonstrated that this distortion can have serious consequences for downstream interpretation, for example, suggesting clusters that do not exist in the original data. Motivated by these developments, we implemented a software package, `distortions`, which builds on state-of-the-art methods for measuring local distortions and displays them in an intuitive and interactive way. Through case studies on simulated and real data, we find that the visualizations can help flag fragmented neighborhoods, support hyperparameter tuning, and enable method selection. We believe that this extra layer of information will help practitioners use nonlinear dimensionality reduction methods more confidently. The package documentation and notebooks reproducing all case studies are available online at https://krisrs1128.github.io/distortions/site/.

## 1 Background

Nonlinear dimensionality reduction methods like UMAP and *t*-SNE are central data visualization tools in modern biology. By projecting high-dimensional molecular profiles into lower dimensions, they reveal salient biological variation across cells. These methods support diverse applications, including developmental trajectory analysis, reference atlas construction, and disease characterization. They are included in widely used data analysis workflows like Scanpy [74] and Seurat [65] and have been popular in practice, reflecting their utility in modern biological research. Nonetheless, these methods have been controversial [7, 29, 37], because they can introduce distortions and artefacts. These shortcomings include exaggerating cluster differences, failing to capture density variation, and suggesting non-existent trajectories [13, 10, 72, 71, 21], which can complicate and cast doubts on the biological interpretation of the observed patterns, potentially leading to false discoveries.

Recent dimensionality reduction methods improve reliability [53, 76, 46, 30]. For example, scAMF [76] constructs nearest neighbor graphs linking cells with their *K* neighbors. scAMF, however, builds these graphs from several transformed versions of the data – rank transformed, *l*^2^-normalized, and log (1 + *x*) transformed. To denoise these transformed measurements, each cell is shrunk towards the average of a local neighborhood derived from the associated graph. Focusing on likely marker genes encourages clearer separation between cell types. This method uses prior knowledge of the relevant transformations and genes to produce more trustworthy visualizations of single-cell RNA expression data.

Recent diagnostics prevent artefacts (such as improved initialization [26] and automatic hyperparameter selection [69, 75, 31]) and to support embedding interpretation, such as adaptations for visualization faithfulness [44, 34] and statistical tests to flag problematic embedding regions [75, 31]. methods. Venna and Kaski (2006) [69] observed a fundamental trade-off between embedding trustworthiness (embedding neighbors reflect original neighbors) and continuity (original neighbors stay together). They offer distortion measures that compare neighbor distances in the original and embedding space, similar to our Equation 3. They aggregate these differences into global trustworthiness and continuity summary statistics, which were used to benchmark existing dimensionality reduction methods and define a new method with a parameter to control the trade-off.

More generally, these approaches produce valuable information, but they fall short of capturing the complexity of nonlinear embedding distortions. Nonlinear distortions are *local, direction-dependent* (*anisotropic*) – stretching in some directions while contracting in others – and *spatially variable*, changing gradually from point to point or abruptly between clusters. This complexity means that global summaries omit key information on local embedding quality. Static visualizations also struggle to faithfully represent distortion context without inducing information overload. Further, existing methods vary in their capacity to remove distortions or provide quantitative measures of the associated improvements. To address these limitations, we introduce the `distortions` package, which addresses the above challenges by building on a mathematically rigorous definition of local metric distortion (RMetric), which is not tied to any particular data embedding algorithm [41], available in our package as `local_distortions()`. Further, since local metrics can change abruptly, we introduce complementary methods for detecting discontinuities (Section 4.6).

Our paper applies RMetric to biological data for the first time, complementing it with a method for flagging fragmented neighborhoods (Section 4.6) that excels when local metrics change abruptly. To render the rich information returned by `local_distortions()`, we introduce a version of the *focus-plus-context principle* [18, 55, 14], supporting the progressive and user-controlled disclosure of sources of distortion based on user interaction, while maintaining the overall query context. This approach helps users interactively flag algorithmic artefacts and answer questions about them that are impossible to answer in full detail with static visualizations. Further, by introducing a new method for interactively isometrizing an embedding, we make it possible to obtain a distortion-free view of the underlying data’s intrinsic geometry in the vicinity of the region of interest.

In summary, this paper makes the following contributions:

1. Applying state-of-the-art measures of local distortion from the manifold learning literature [52, 40, 41] to single-cell data for the first time. Our package is the first to quantitatively measure and render the anisotropy of the local distortions, along with the points of local embedding discontinuity.
2. Demonstrating the practical utility of distortion measures in choosing between algorithms and hyperparameters. We find that these metrics can reveal systematic differences in the interpretation of embedding distances across cell types and highlight contiguous neighborhoods that become fragmented during dimensionality reduction. Thus, they support objective comparison of embedding results, and the accompanying visualizations provide insight into qualitatively different types of distortion.
3. Developing interactive visualizations that highlight distorted regions and enable local corrections. We introduce an isometrization method that allows users to interactively correct distortions locally within regions of interest. Additional focus-plus-context approaches reveal distorted neighborhoods based on user queries of summary visualizations.

We validate this functionality using data with known low-dimensional structure, then apply the package to three single-cell datasets, showing the potential for improved biological interpretation and nonlinear embedding method application. The package is hosted at https://pypi.org/project/distortions/ and documented at https://krisrs1128.github.io/distortions/site/.

### 1.1 Distortion estimation

To set up our results, we briefly review distortion estimation. Embedding methods aim to learn a lowdimensional, potentially nonlinear manifold on which the data lie. This manifold hypothesis is motivated by the fact that only certain patterns of gene expression are plausible, due to regulatory constraints. Geometrically, every point on the manifold can be mapped to a local coordinate system, called a *chart*. Biologically, the local coordinates are directions of shifting activity of latent biological processes. An ideal embedding method would perfectly recover these intrinsic charts, ensuring that distances on the biological manifold ℳ are reflected in the embedding. Such a distance-preserving manifold embedding is called an *isometry*.

Even in linear dimensionality reduction, distances require careful interpretation. For example, in principal component analysis (PCA) plots, it is recommended that the axes be rescaled to reflect the relative variances explained by each component [45]. This issue becomes more difficult in nonlinear settings, where the interpretation of relative distances can vary locally across regions of the visualization [52]. Practical algorithms inevitably introduce distortion, systematically dilating some directions while compressing others. Depending on the direction of travel and the starting point, the same distance in the embedding space might correspond to different distances along the manifold. Though we may not be able to avoid distortion, we can at least estimate it. Here we will call this estimate RMetric (Section 4.3 explains this name) and can be represented in various equivalent ways, as shown in Fig 1. For instance, the function `local_distortions()` returns RMetric as a matrix **H**^(*i*)^. The matrix **H**^(*i*)^ gives a quantitative measure of the distortion induced locally around data point *i* by an embedding method. A mathematical treatment is provided in the Methods section, and we refer to [42] for an in-depth discussion.

**Fig. 1:**
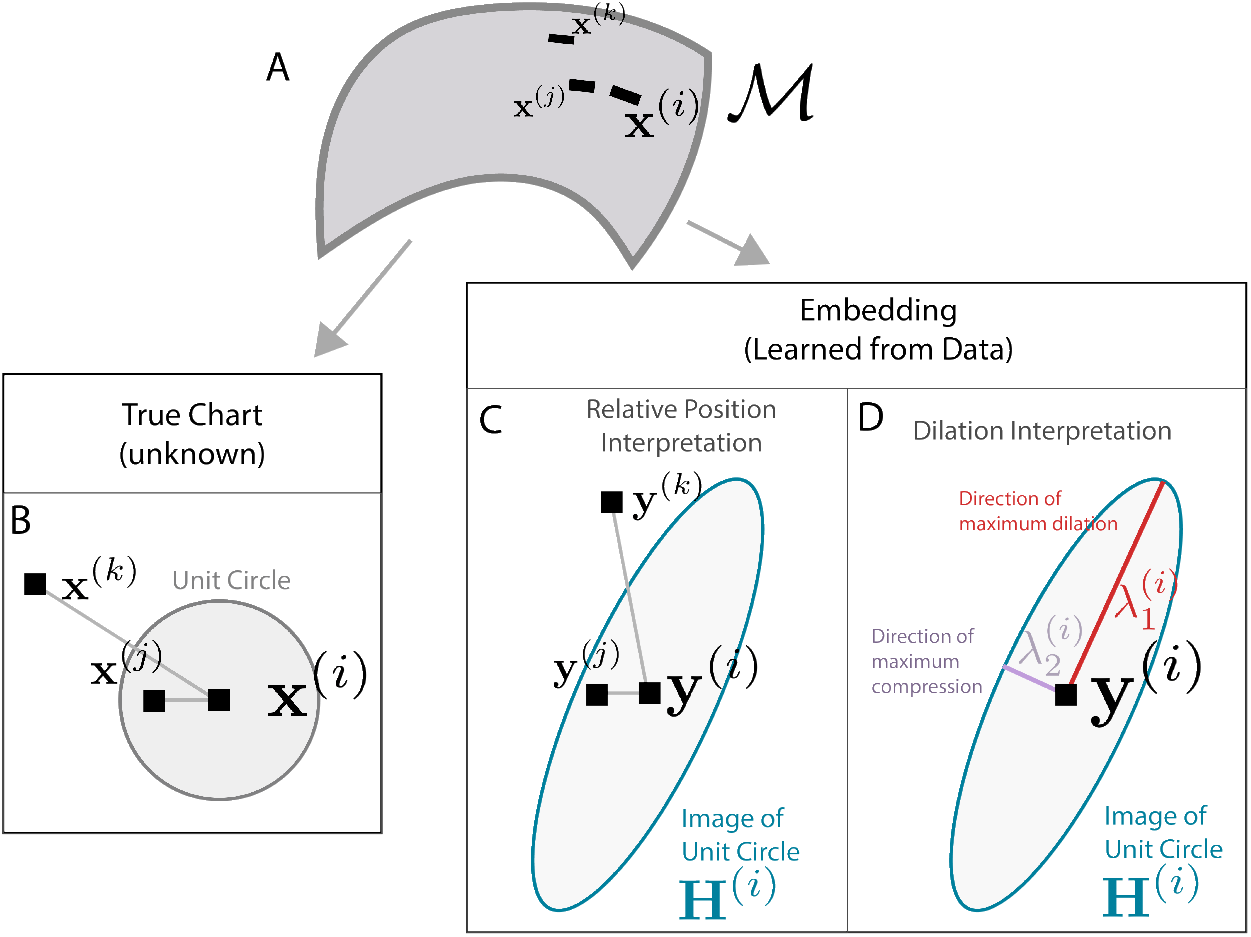
Interpreting the matrices **H**^(*i*)^ generated by the RMetric algorithm. A. Three points on a hypothetical manifold ℳ. B. The three points from panel A arranged on one of the charts that defines ℳ. A unit circle with respect to the intrinsic metric around ***x***^(*i*)^ is overlaid. C. The embedding algorithm distorts the unit circle from A. Though the distances and angles between samples have changed, the ratios of their distances to the unit circle have not. Though the true distortion around ***y***^(*i*)^ is unknown, it can be estimated using **H**^(*i*)^ (blue ellipse). D. The same **H**^(*i*)^ as panel C, but emphasizing the directions and degree of maximum dilation and compression.

We visually encode the local distortions **H**^(*i*)^ with ellipses, displayed at each embedded point. Ellipses with circular shapes reflect regions where the embedding approximates an isometry. Thus the size and orientation of ellipses gives the principal directions of stretch/compression around point *i*, and the ellipse itself can be seen as a polar plot of the stretch (or compression) associated to each direction from point *i* (Fig 1). Specifically, larger ellipses appear when distances have been inflated and the major axes appear in the direction of most extreme dilation. This approach generalizes Tissot’s indicatrix from cartography [27] to high-dimensional embedding algorithms.

## 2 Results

### 2.1 Detecting cluster-specific differences in local metrics

This section gives two examples where local metric visualization highlights systematic differences in embedding interpretation across clusters.

#### 2.1.1 Gaussian mixtures with different variances

We evaluate the recovery of intrinsic geometry in an embedding of a mixture of two Gaussians. We sampled 500 points each from two components: 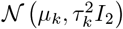 where *µ*_*A*_ = (0, 0)^⊤^, *µ*_*B*_ = (30, 0)^⊤^, *τ*_*A*_ = 10, and *τ*_*B*_ = 1. The resulting mixture is shown in Fig 2A. We applied UMAP with 50 neighbors and a minimum distance of 0.5. Despite the large differences in variance, UMAP returned clusters with comparable sizes and densities (Fig 2B). We applied the RMetric algorithm with a geometric graph Laplacian constructed from the 50-nearest neighbor graph and rescaling *ϵ* = 1. The affinity kernel radius was set to the mean of the original data distances between neighbors on this graph. We obtained RMetric singular values 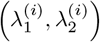 using a singular value decomposition of the dual of the embedding Riemannian metric, as detailed in Section 4.3. To prevent samples with outlying RMetric singular values from obscuring variation among the remaining points, we truncated *λ*^(*i*)^ at a maximum value of 5; this affects 6 samples. To ensure that isometrization does not uniformly contract or expand neighborhoods across the visualization, we further divided all **H**^(*i*)^ by a scaling factor 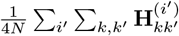.

**Fig. 2:**
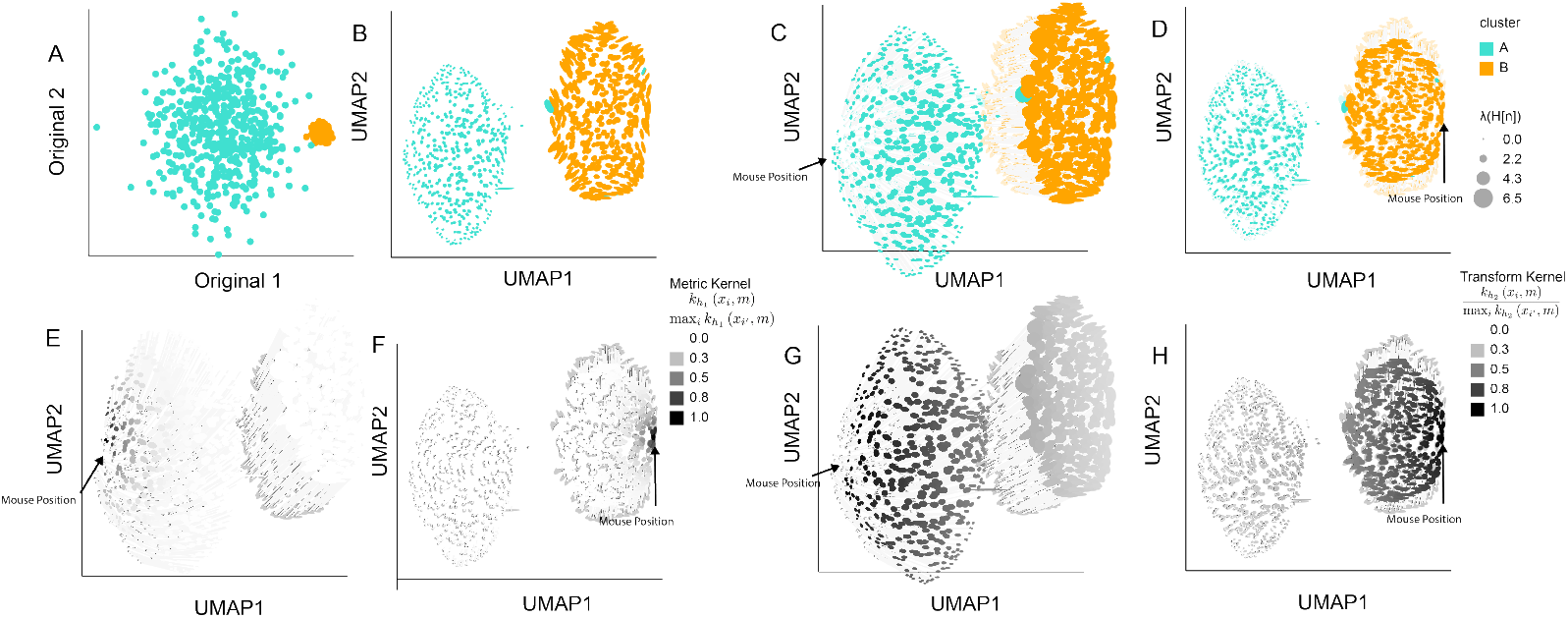
Interactive isometrization partially restores density differences in an Gaussian mixture embedding. A. Original simulated data. Cluster B has smaller variance compared to Cluster A. B. Ellipse orientation and sizes encode differences in local metrics in the UMAP embedding. Smaller ellipses mean that the same distance in the embedding space corresponds to larger distances in the original data space. C. The isometrization interaction updates ellipse size and positions to reflect the local metric in the hovered-over region. This partially restores the difference between cluster variances that were lost in the initial embedding. D. The analogous isometrization when hovering over Cluster B (orange). Cluster B slightly shrinks, while Cluster A (blue) remains at its original size. E. The normalized kernel similarities defining the contribution of each **H**^(*i*)^ to the **H**^∗^ used in the isometrization from panel B, as given in equation (4). F. The analog of panel E for the mouse interaction in Panel D. G. The normalized kernel similarities describing the extent to which each point is moved from its original position, as given in equation (5). The analog of panel G for the mouse interaction in panel D.

The resulting local metrics **H**^(*i*)^ are overlaid as ellipses in Fig 2B-H. Fig 2B shows that Cluster *A* has smaller ellipses than Cluster *B*, correctly reflecting the differences in cluster variance lost by the UMAP embedding. Fig 3 shows the coordinates of the truncated *λ*^(*i*)^ plotted against one another. The clear separation in singular values across clusters reinforces the qualitative differences in ellipse sizes from Fig 2A. Fig 2C-D show the isometrized versions of Fig 2B when hovering over samples in Cluster A and B, respectively. These interactions recalculate the embedding locations and ellipse sizes to bring the local metrics **H**^(*i*)^ near the viewer’s mouse position closer to the identity *I*_2_, resulting in more circular ellipses. Thin grey lines connect the isometrized and the original embedding coordinates. When hovering over Cluster A, the samples in that cluster become spread further apart, while those in Cluster B are translated to the right but remain at their original density. In contrast, when hovering over Cluster B, the samples in that cluster contract while those in Cluster A remain close to their original positions.

**Fig. 3:**
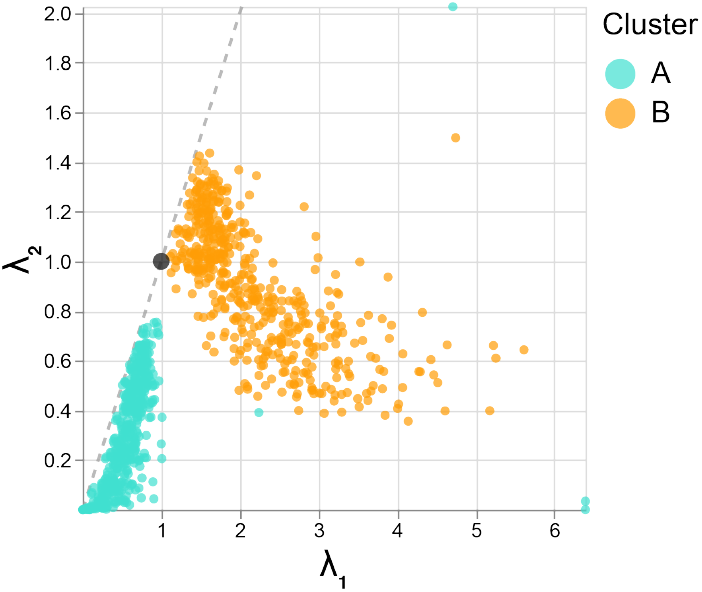
Singular values 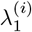 and 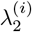 of the **H**^(*i*)^ estimated in Figure 2. The larger 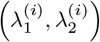 in Cluster B results in the larger ellipse sizes for that cluster, indicating that embedding distances for this cluster have been “spread out” relative to original, pre-embedding distances. This effect is consistent with the data shown in Figure 2A.

More precisely, Fig 2C calculates a “local” metric **H**^∗^ based on the weights in Fig 2E, which are high (darker) near the viewer’s mouse position. An exact isometrization with respect to the current region of interest would update embedding coordinates ***y***^(*i*)^ to 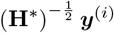 [52] across the entire visualization. We instead restrict the transformation to areas close to the viewer’s current interaction region. Informally, the darker points in Fig 2G are allowed to be updated more aggressively than the lighter points; the formal transformation is detailed in equation (5). The analogs of Fig 2E and G for the interaction in Fig 2D are given in Fig 2F and H. Comparison with additional distortion visualization techniques is given in Supplementary Section A.3.

#### 2.1.2 Local metrics vary across cell types in a PBMC atlas

We next analyze peripheral mononuclear blood cell (PBMC) single-cell genomics data (2683 cells, 1838 genes) from 10X Genomics, with default processing from scanpy [60, 74], which included total sum scaling (TSS), a log (1 + *x*) transformation, and highly variable gene filtering. We denoised the data with PCA, using the default 50 components from `scanpy.pp.pca`. Following the workflow at [59], we apply UMAP with 10 neighbors and minimum distance set to 0.5. Cell types were identified with Leiden clustering and canonical marker genes CD79A and MS4A1 (B cells), FCER1A and CST3 (dendritic cells), GNLY and NKG7 (NK cells), FCGR3A (monocytes), IGJ (plasma cells), and CD3D (T cells). To estimate the data’s intrinsic geometry, we applied RMetric using the geometric graph Laplacian constructed from the 10-nearest neighbor graph and rescaling *ϵ* = 5. The affinity kernel radius was set to three times the mean of the original data distances within this graph. To prevent a few highly skewed ellipses from influencing the remaining points, we truncated the singular values 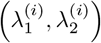. from above at 2.5. We divide by the same 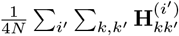 scaling factor as in Gaussian mixture example above.

The resulting ellipse-enriched embedding Fig 4A reveals systematic metric differences across cell types. T cells ellipses are oriented with major axes in the northwest/southeast direction, suggesting that distances orthogonal to this direction compressed in the embedding. In contrast, dendritic cells are generally oriented in the southwest/northeast direction, suggesting greater spread away from the monocytes than the embedding alone indicates. Fig 4D displays the truncated and rescaled singular values 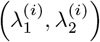. Points closer to the *x*-axis correspond to ellipses that are more eccentric than those near the center of the plot. Cell types differ systematically in this view as well, reinforcing our conclusion that local metrics are associated with cell type. The panel also draws attention to the high condition numbers among subsets of the T and NK cells. In contrast, many monocytes lie in the middle of the panel; these are the more circular embeddings in Fig 4A.

**Fig. 4:**
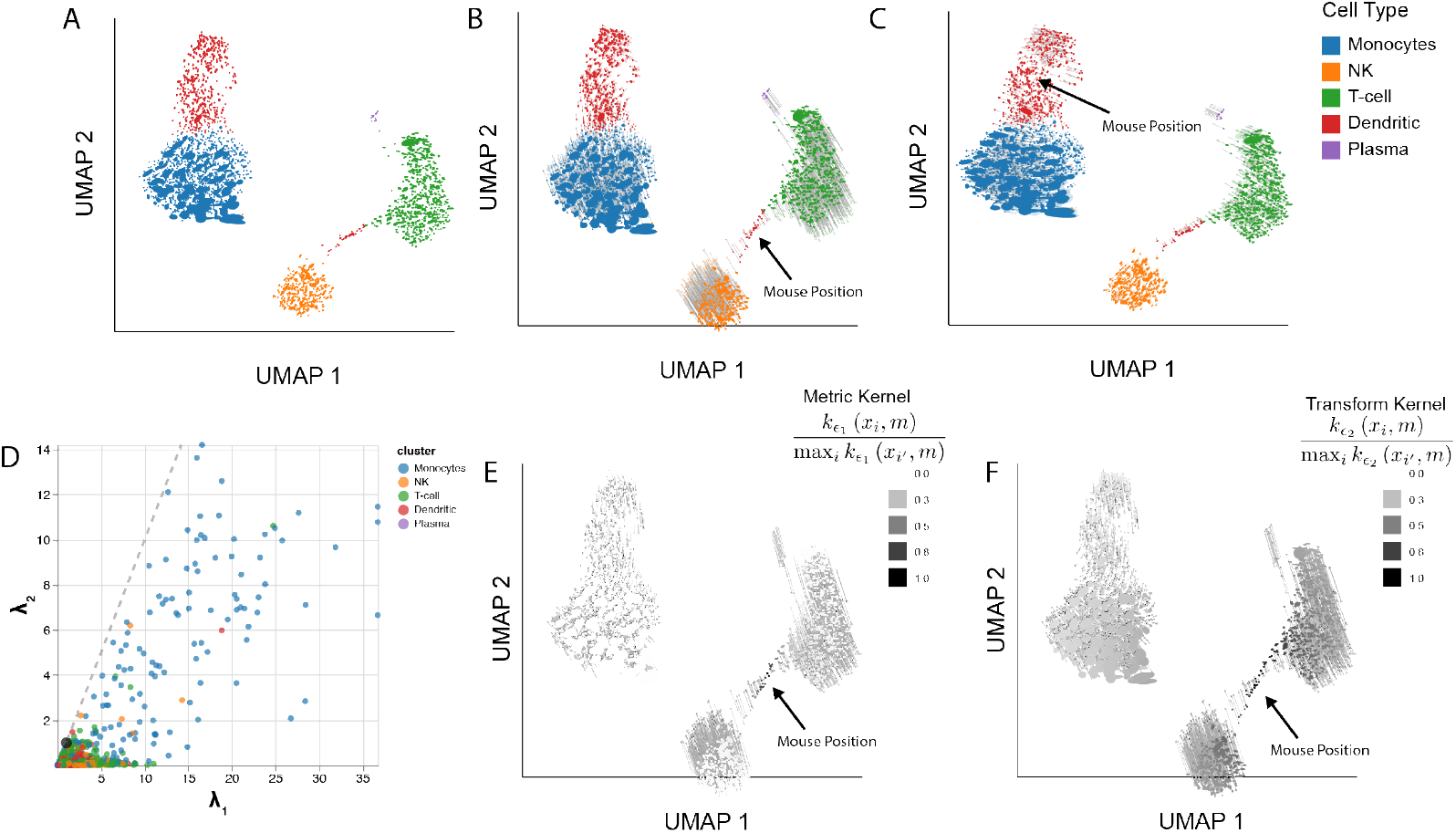
Isometrization of the PBMC UMAP embeddings. A. Ellipse orientation and sizes vary systematically across regions of the embedding, indicating differences in local metrics within and between cell types. B. An updated version of panel A when the mouse is positioned over a subset of dendritic cells that bridge the NK and T cell clusters. Transparent ellipses mark the cells’ original positions, and lines connect the original and updated locations. C. Isometrization when hovering over dendritic cells. D. The windsorized singular values 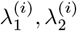 associated with **H**^(*i*)^ across cells. Ellipse size is determined by 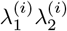 and eccentricity by 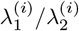. A version that zooms into the region near the origin is given in Appendix Fig 16. E. The normalized kernel similarities defining the local metric **H**^∗^ (equation (4)) when the mouse is placed as in panel C. F. The analog of panel E when the mouse is placed as in panel D.

The slopes in Figure 4D have a qualitative meaning, reflecting the typical ratio 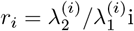, with values near one indicating nearly isotropic local geometry and smaller slopes correspond to stronger directional anisotropy, where the embedding stretches on direction in the underlying manifold is more than another. The product 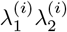 reflects the local volume change induced by the embedding. Points farther from the origin lie in regions that are locally expanded, while those closer to the origin lie in locally compressed regions. Further, zooming into Fig 4D, Appendix Fig 16 reveals a second monocyte subset with smaller singular values. This pattern matches the bimodality in monocyte size distribution in Figure 4A. The UMAP embedding appears to have collapsed the two monocyte subsets, compressing distances for smaller points and dilating them for larger points. Though changing the visual markers from circles to ellipses is a small difference, the associated local metrics reveals valuable context about how UMAP warps intrinsic geometry across the visualization.

Fig 4 illustrates isometrizations for two cell types. Fig 4B - C show the embedding after placing the cursor over clusters of dendritic cells (Fig 4B -C) In both panels, the solid ellipses represent the updated embedding, while transparent ellipses and thin lines indicate the original positions and metrics. Isometrization over the dendritic cells in panel B expands the main dendritic cluster and increases the distance from the NK cluster and T cell clusters. Figs 4E - F display the normalized kernel similarities used to define the local metric **H**^∗^ and the regions of transformation. The interaction over the second group of dendritic cells (Fig 4C) increases the spread of cells close to the mouse position. Monocyte orientations are shifted slightly, but other cell types remain largely unchanged. Together, this suggests that the distortion of first group of dendritic cells is more severe in this choice of UMAP embedding. Finally, additional baseline comparison is given in Supplementary Section A.3.

#### 2.2 Identifying fragmented neighborhoods

There is emerging evidence that nonlinear dimensionality reduction methods can introduce embedding discontinuities [31], meaning that some points that are nearby in the original space end up being embedded as far from one another as those that are originally very different. In particular, points that lie in the same neighborhood in the original space may be fragmented into different embedding regions, complicating the interpretation of between-cluster relationships. To address this, `distortions` provides metrics for quantifying fragmentation at the neighborhood and pair levels, based on the relationship between observed vs. embedding distances among nearby points in the original data space, as detailed in the Methods (“Identifying fragmented neighborhoods”). A focus-plus-context visualization approach [18, 28, 55] then allows viewer interactions to progressively reveal the extent of fragmentation within different embedding regions. We provide examples below.

##### 2.2.1 Mammoth skeleton

We evaluate this strategy on UMAP embeddings of a three-dimensional mammoth skeleton point cloud (Fig 5A) generated by the Smithsonian Museums in an effort to digitize their collection [47]. This dataset has been used to study the artifacts introduced by UMAP [10]. It has the advantage of being directly visualizable in three dimensions (Fig 5A). Further, the data exhibit patterns at both global and local scales. For example, a successful dimensionality reduction method must preserve global relationships, like the relative positions of tusk, skull, and legs, and also fine-scale differences, like the distinction between bones in the rib cage. We applied UMAP (50 neighbors, minimum distance 0.5) to embed the 10,000 samples available in these data. The RMetric algorithm was applied using a geometric graph Laplacian constructed from the 50-nearest neighbor graph and a rescaling *ϵ* = 5. The affinity kernel radius was set to 3 times the mean of the original data distances between neighbors on this graph. The resulting **H**^(*i*)^ are directly encoded using ellipse dimensions without any post-processing. To identify distorted pairs, we used the boxplot display with outlier threshold set to 10×IQR. To identify distorted neighborhoods, we applied the bin-based screening metric (see Methods) with *κ* = 0.1 and *σ* = 3, requiring that at least 10% of neighbor distances be poorly preserved. This flags 425 potentially fragmented neighborhoods.

**Fig. 5:**
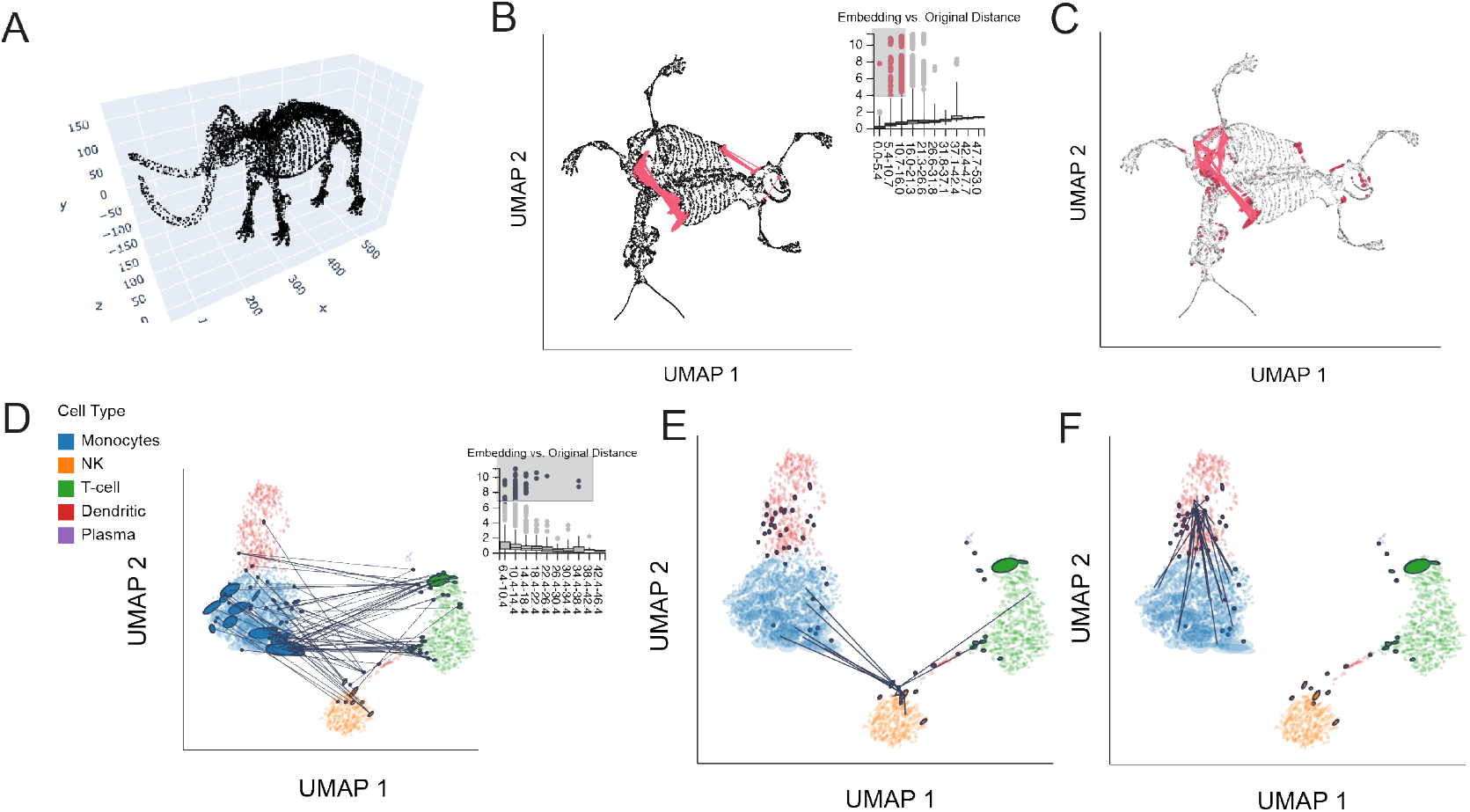
Fragmented neighborhoods and links. A. The original mammoth point cloud, before applying any dimensionality reduction. B. Pairs with poorly preserved distances in the mammoth data. The viewer has selected pairs of points that are close to one another in the original space, but which are far apart in the embedding. C. Analogous poorly fragmented neighborhoods defined using the quantile smoothing criteria. D. Pairs with poorly preserved distances in the PBMC data. Distances between NK cells, T cells, and monocytes have been exaggerated by the UMAP embedding. E. A subgroup of NK cells with poorly preserved neighborhoods in the PBMC data. Cells close to the viewer’s mouse interaction have neighbors spread across T cells, NK cells, and monocytes. F. A different subgroup of NK cells with fragmented neighborhoods. These cells often have neighbors lying on the far boundary of monocytes.

Fig 5B shows the boxplot widget overlaid on the UMAP. Reassuringly, the median embedding distances increase monotonically as the distances in the original space increase. However, within each bin, the distribution of embedding distances is skewed, especially for small distances in the original data space. Many pairs of points within these bins appear much further apart in the embedding than expected. In the current display, the viewer has selected the outliers within the three leftmost bins, highlighted in pink. The corresponding pairs are linked together in the main embedding view. These pairs include points on the left and right shoulders of the mammoth. These points are close to one another in three dimensions, but have been spread apart by the embedding. UMAP appears to reflect geodesic rather than Euclidean distance, effectively “flattening” the mammoth skeleton. In addition to the left and right shoulder pairs, the highlighted outliers include neighbors where one point lies on the last right-side rib bones and the other on the right side of the pelvis. UMAP embeds these adjacent bones further apart than appropriate, another distortion of the original structure.

The fragmented neighborhoods displays in Fig 5C confirms these findings. For example, the flattening of the shoulder is evident in the chain of fragmented neighborhoods in this region. Points further along the rib on the right and pelvis are also highlighted, as in the boxplot view. In this case, the viewer’s mouse lies over the right shoulder of the mammoth. Unlike the boxplot view, this allows us to view all the neighbors of distorted points near the mouse, showing the neighbors along the chest and arm whose distances are not outlying in the boxplot. This view also reveals more isolated fragmentations, for example on the skull, arms, and tail. Hovering over these points shows that a large fraction of their neighbors have also been spread apart (e.g., left and right hand sides of the skull), even if their absolute embedding distances are not large enough to stand out in the boxplot interactions.

##### 2.2.2 PBMC gene expression

We next identify distorted embedding pairs in the PBMC example. The boxplot widget again reveals outliers with exaggerated embedding distances (Fig 5D). The viewer has brushed the top outliers across all bins. This selection highlights pairs of T cells and monocytes that are close despite the apparent embedding separation. This view distinguishes between two regions with distorted pairs within the T cell cluster. Unlike the dendritic cells, which visibly cluster into two subtypes, these two subtypes of distorted T cells do not stand out from the main T cell cluster. Nonetheless, both subtypes are near the boundary (top and bottom left, respectively) of the overall cluster. This suggests that in high dimensions, the T cell cluster may be curved in a way that allows these subgroup to be closer to the monocytes than is visible in the embedding.

We next identify fragmented neighborhoods using the bin-based strategy (*L* = 10, *κ* = 0.2, *σ* = 2 threshold), resulting in 72 cells with fragmented neighborhoods. Fig 5E highlights fragmented NK cell neighborhoods that are are connected to both T cells and monocytes. These cells appear to bridge several cell types. Fig 5F shows a subgroup of distorted dendritic cells. Hovering over them shows that they are neighbors with distant monocytes. Two subtypes of T cells are flagged as distorted; these largely overlap with those highlighted by the boxplot visualization in Fig 5D. Only eleven monocytes with fragmented neighborhoods are flagged, suggesting that this cluster does not suffer from fragmentation as severely as the others. Interactive distortion visualization can reveal different degrees and types of distortion across and within cell types.

#### 2.3 Guiding method selection and tuning

In addition to interpreting individual embedding visualizations, distortion metrics can be used to compare different embedding methods and hyperparameter choices. They give a quantitative way to judge how well competing embeddings preserve the original data’s structure. Further, the interactivity implemented in `distortions` makes it possible to explore where distortions arise, without overwhelming viewers with all contextual information at once. In this section we use three example datasets to illustrate how distortion visualization can guide method selection and tuning.

##### 2.3.1 Clarifying how hyperparameter choice impacts distortion

Hyperparameters in nonlinear dimensionality reduction methods like UMAP and *t*-SNE can substantially influence results [26, 5, 72]. Distortion visualization can reveal the trade-offs imposed by specific choices. We evaluate this using the hydra cellular differentiation data from [63]. This study used single-cell RNA sequencing to measure gene expression of a developing hydra polyp, an organism notable for its regenerative ability. Fig 1 of their paper is a *t*-SNE that clarifies the cellular composition of hydra tissue as well as the differentiation paths from stem and progenitor cells to specialized cell types. To create a setting with greater statistical instability and where hyperparameters may play a more important role, we take a random sample of 2000 of the original 24,985 cells, though see Supplementary Section A.1 for further discussion on scalability. As in the analysis of the PBMC data, we apply TSS normalization, a log (1 + *x*) transformation, and filter to the top 1000 highly variable genes.

Following the supplementary analysis of [63], we apply *t*-SNE to PCA denoised data (top 31 components). We compare results when using perplexity values of either 80 or 500. To estimate distortion we use the RMetric algorithm with a geometric graph Laplacian with 50 nearest neighbors and a rescaling *ϵ* = 5. The radius for the affinity kernel was set to three times the average original data distance in the 50-nearest neighbor graph. Both the boxplot widget and the neighborhood fragmentation visualizations suggest qualitatively different types of distortion across the two perplexity settings. At a perplexity of 80, the fragmented neighborhoods occur in the gaps between cell type clusters (Fig 6A). Hovering over these neighborhoods reveals connections to adjacent cell types (Fig 6C), suggesting that some transitions in gene expression programs between cell types may in fact be more gradual. These blurrier transitions are captured at a perplexity of 500 (Fig 6B). However, at this hyperparameter value, many fragmented neighborhoods appear along the top and bottom boundaries of the embedding. Interacting with the display reveals that at this hyperparameter choice, the embedding fails to preserve distances between peripheral neighborhoods. For example, the viewer’s selection in Fig 6D highlights neighbors that have been split across opposite sides of the visualization.

**Fig. 6:**
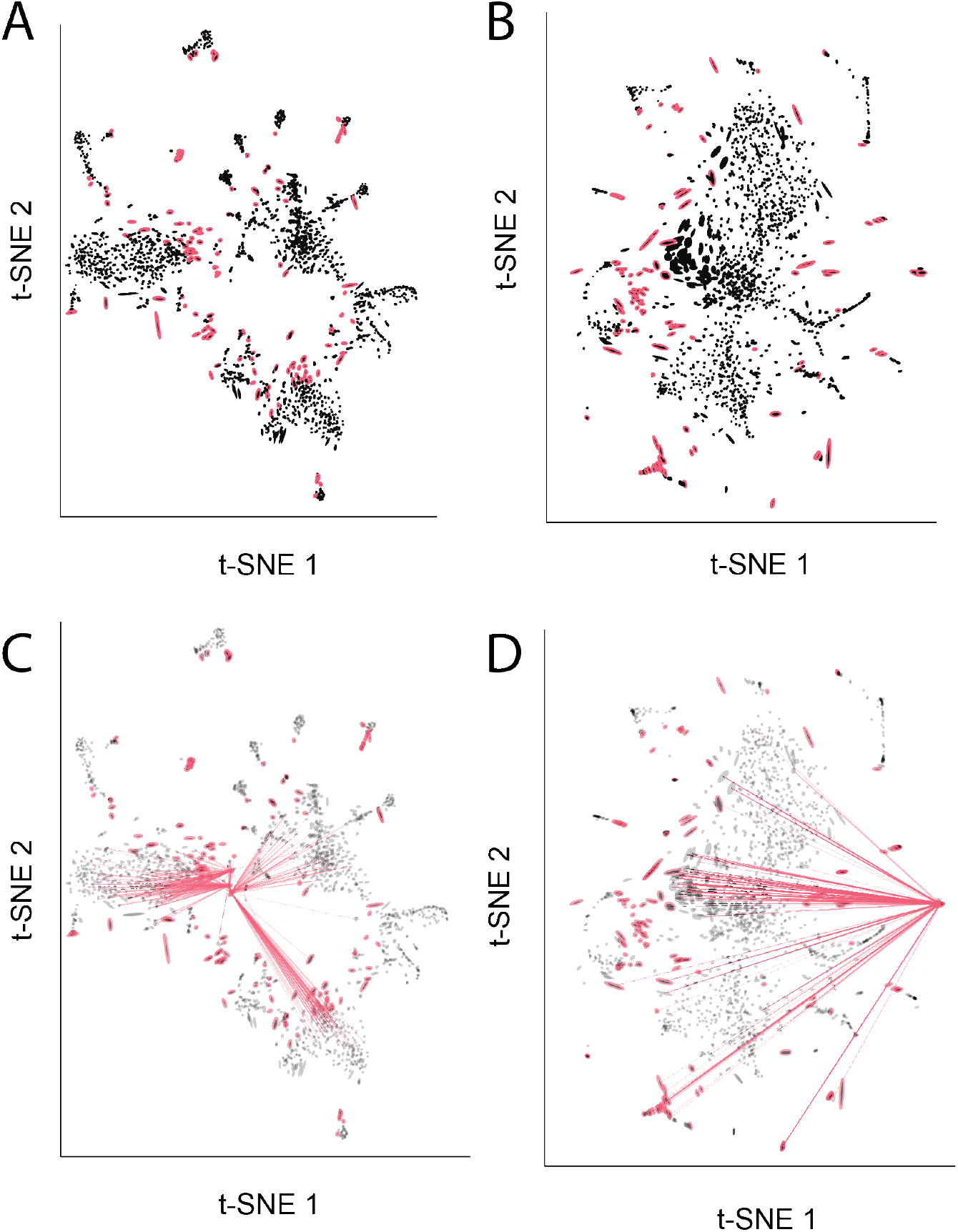
Salient characteristics of distortion vary across hyperparameter settings. A. The *t*-SNE embedding of the hydra cell atlas dataset when perplexity hyperparameter is set to 80. This embedding exaggerates the distinction between cell type clusters. B. The analogous view when the *t*-SNE perplexity is set to 500. At this hyperparameter value, the main clusters are now more overlapping, but the distances along the periphery of the embedding are less well preserved. C. Hovering over fragmented neighborhoods near the bottom-left of the embedding in panel A shows that neighbors are often shared between clusters. D. Hovering over a fragmented neighborhood in Panel B shows that points near the periphery can be neighbors with points spread throughout the visualization.

This reveals a basic trade-off: higher perplexity better reflects distances between main cell types but arbitrarily places rarer types, while lower perplexity correctly places these rare clusters correctly at the cost of inflating distances between common cell types. This additional context gives confidence in the conclusions drawn within specific regions of separate visualizations. These conclusions can still be reliable even when no single view preserves all relevant properties of the original high-dimensional data. Further, though the qualitative differences between hyperparameter choices would be difficult to obtain through manual inspection of the distances within the embedding output, the interactive display allows the differences to pop out naturally.

##### 2.3.2 Comparing initialization strategies using distortion metrics

Nonlinear dimensionality reduction methods can be sensitive to initialization strategies. Indeed, most single cell analysis packages use a preliminary dimensionality reduction step, like PCA or Laplacian eigenmaps [3], to initialize the optimization [26, 57]. We next study whether distortion metrics can detect issues arising due to poor initialization. To this end, we rerun the UMAP analysis of the PBMC data and consider a random, rather than the default spectral, initialization. All other dimensionality reduction and visualization hyperparameters remain as before. Fig 7 presents the results. Compared to Fig 4A, Fig 7A separates the NK cells into distinct groups falling on opposite sides of a main cluster of dendritic cells and monocytes. Brushing outlying neighbor distance pairs in the boxplot in Fig 7C highlights the fact that these two groups share many neighbors, and that the gap is artificial: many NK cells are neighbors with T cells despite lying on opposite sides of the plot. This suggests that the spectral initialization, which places T cells and NK cells adjacent to one another, better preserves their neighborhood relationships.

**Fig. 7:**
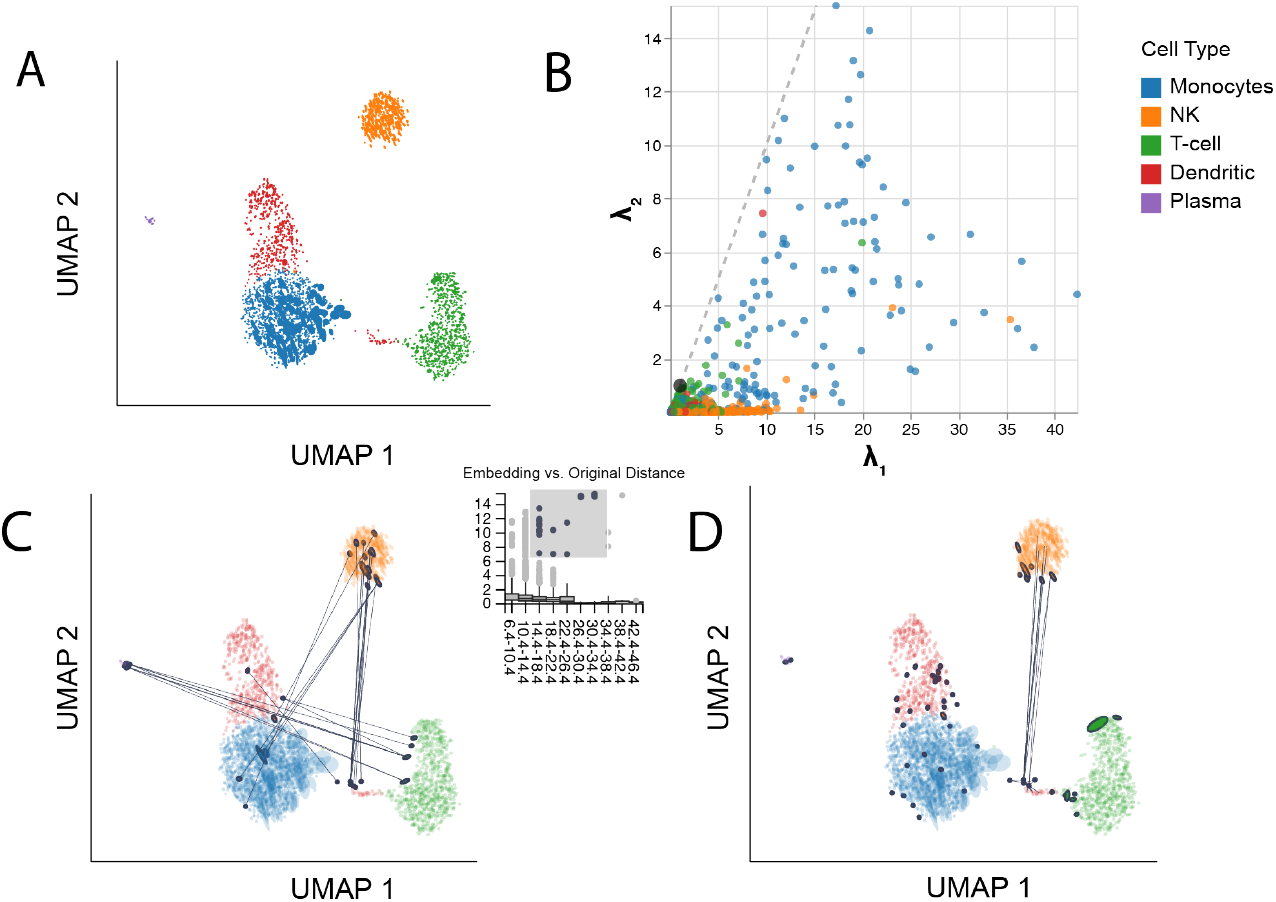
Distortion visualizations highlight problems with randomly initialized UMAP. A. UMAP embedding of the PBMC data when applying random initialization. B. The analog of Fig 4D in the random initialization setting. The systematically larger condition numbers 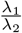 correspond to more eccentric ellipses in panel A. C. Brushing over pairs with large embedding vs. original distances highlights Dendritic-NK and Plasma-T cell neighbors whose relative distances are poorly preserved. These cell types are placed close to one another in the spectral initialization of Fig 4. D. Hovering over the fragmented neighborhoods in the bottom right corner of the plot highlights dendritic cells with more neighbors among NK cells, despite their close placement to T and monocyte cells. This subtype of dendritic is placed close to the NK and T cells in Fig 4.

Fig 7D displays the fragmented neighborhoods, analogous to Fig 5E. Though some T-cell-adjacent NK cells had been flagged in the spectrally-initialized embedding, a larger number are distorted in the random initialization, including many with neighbors in the monocyte cluster. Further, the reduced *y*-axis range in Fig 7B relative to Fig 4D draws attention the greater eccentricity of ellipses in the random initialization, indicating larger distortion of local metrics. Importantly, none of these issues with the random initialization are detectable from the embedding coordinates alone. Both ellipse eccentricity and interaction with distortion summary metrics add context for understanding the importance of effective UMAP initialization.

##### 2.3.3 Analyzing density preservation in a *Caenorhabditis elegans* cell atlas

We applied our package to a single-cell atlas of *Caenorhabditis elegans* development [49], originally gathered to characterize the gene programs activated during different phases of embryogenesis in the *C. elegans* model system. These data include measurements on 86,024 cells, of which 93% have been manually annotated with cell types by the authors. Nonlinear embeddings applied to this dataset are known to obscure meaningful differences in local density, causing biologically meaningful cell types to appear sparser or denser than appropriate [44]. Therefore, we compared UMAP with the density-preserving algorithm DensMAP and use `distortions` to evaluate the improvement in local metric preservation. By utilizing our package’s distortion summaries, we can highlight neighborhoods that are artificially fragmented in the embeddings and quantify the reduction in distortion made possible through the DensMAP algorithm.

Following [44], we consider a PCA-denoised version of the data (top 100 dimensions). We applied both UMAP and DensMAP with 10 neighbors and a minimum distance of 0.5. To simplify the distortion analysis, we considered a random sample of 5000 cell embeddings from each of 10 representative cell types from across the main neuronal, endoderm, mesoderm, and ectoderm lineages in *C. elegans* (ciliated amphid neuron, pharyngeal neuron, GLR, glia, intestine, body wall muscle, hypdermis cell types). This restriction is analogous to focusing on a subset of cell types when testing whether putative cell types are truly distinct or a visualization artifact [64, 15], and it is necessary for avoiding overplotting, an issue discussed further in Supplementary Section A.1. We used RMetric to estimate local metric distortion using a geometric graph Laplacian based on the 10-nearest neighbor graph and affinity kernel radius set to three times the average original neighbor distance in this graph. To identify distorted neighborhoods associated with each method, we apply the bin-based strategy with *L* = 10, *κ* = 0.4, *σ* = 3 to flag points where a fraction of at least 40% of neighbors have embedding distance at least 3 × IQR away from the median within the corresponding bin of original distances.

Fig 8A-B shows the resulting fragmented neighborhoods. In both embeddings, the degree of fragmentation varies by cell type. For example, pharyngeal neuron neighborhoods are often fragmented by both algorithms, while few fragmented neighborhoods are centered on hypodermis cells. Qualitatively, the DensMAP embedding is less compressed into tight clusters than UMAP, suggesting that UMAP may artificially inflate the embedding space densities. Despite using the same graphical encoding scales, the UMAP ellipses also appear to be less uniformly sized. The more compact “hair” plots reinforce this conclusion (Fig 8D-E). Each segment corresponds to one ellipse in panels A - B. The segments are oriented along the minor axis of the ellipses, and their lengths encode condition number 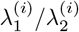. We note that these hair-like graphical marks can be substituted for ellipses in all visualizations and interactions discussed above, including the boxplot and isometrization displays.

**Fig. 8:**
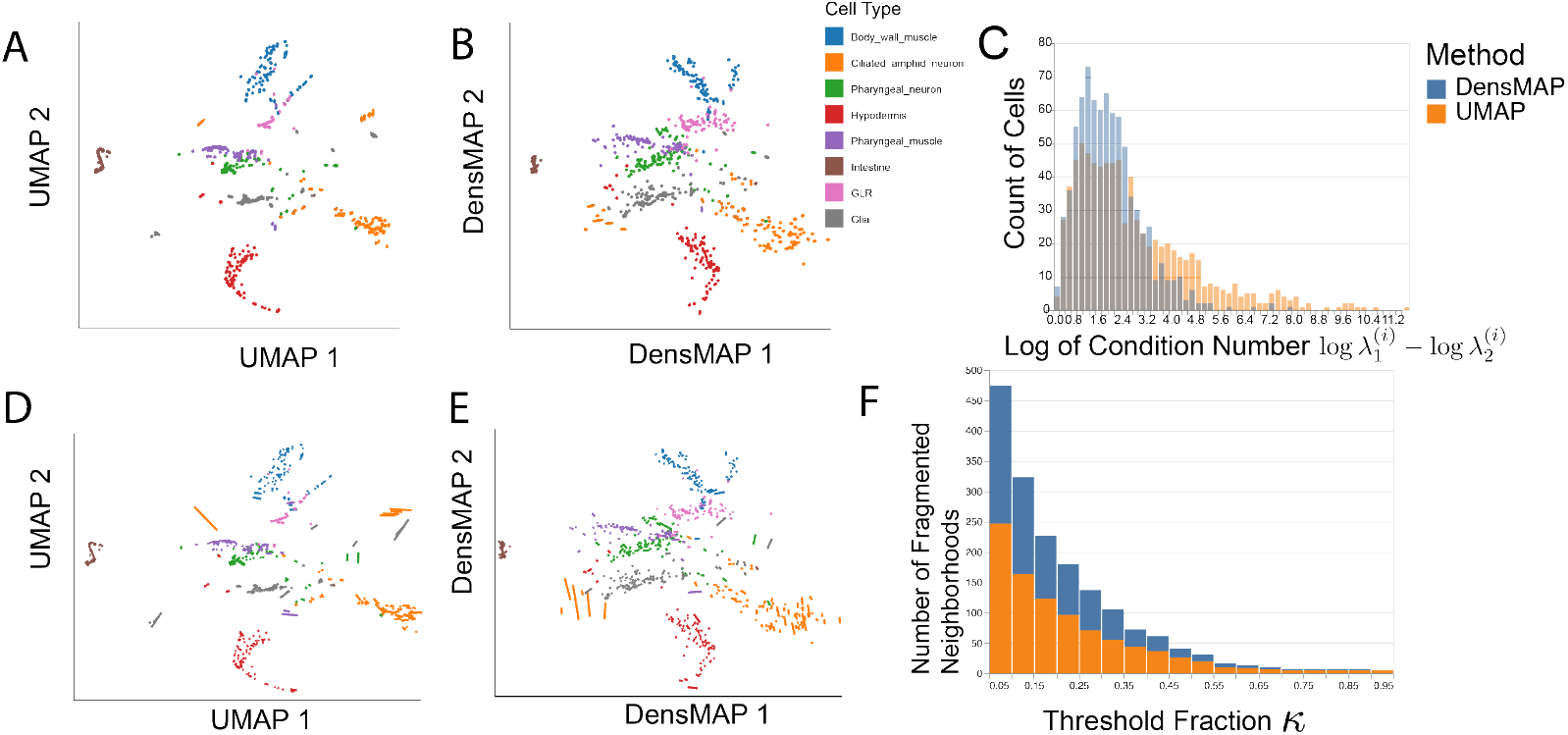
Distortion metrics support comparison of UMAP and DensMAP embeddings. A. A UMAP embedding of the *C. elegans* dataset, subsampled as described in the main text. Ellipses represent the local metric **H**^(*i*)^ across observations. B. The analogous DensMAP visualization. C. Distribution of the (log-)condition numbers 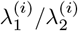 for the UMAP and DensMAP embeddings. Note that the two colors are semi-transparent and partially overlap in the range of low-condition numbers. A value of 0 indicates that the dilation/contraction is the same in all directions, while ce larger condition numbers correspond to more extreme eccentricity; hence this view indicates that DensMAP is more isotropic. D. The analog of panel A using “hair” graphical marks in place of ellipses, to reduce overplotting. The orientation of each segment is orthogonal to the ellipses’ major axes, and the length encodes the condition numbers 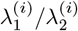. The analogous “hair” plot of panel B. F. A stacked barplot of the number of fragmented neighborhoods when applying DensMAP and UMAP when varying the threshold parameter *κ* used in neighborhood definition. Across a range of thresholds, DensMAP results in fewer fragmented neighborhoods compared to UMAP.

Further, the distortion metrics provide quantitative support of DensMAP’s superior ability to preserve intrinsic geometric information. For example, the histogram in Fig 8C shows that the UMAP resulted in systematically larger metric condition numbers, suggesting more systematic metric distortion. Further, Fig 8F shows that across choices of *κ*, the DensMAP results in fewer fragmented neighborhoods than UMAP. In this case, the distortion metrics led to a stable conclusion across embedding algorithm hyperparameters.

But it is also worth considering the variation of distortion estimates themselves, with respect to their own hyperparameters. In the case of fragmented neighborhoods, it is easy to see that, whether a neighborhood is flagged as fragmented can be dependent on the choices (*L, κ, σ*). There is no specific rule for choosing these, as they should reflect the user preferences.

This is not the case for the local distortion estimates **H**^(*i*)^ and we study their dependence on the hyperparameters in the next section.

#### 2.4 Robustness of RMetric across hyperparameters

For the RMetric the radius and scaling parameters have optimal values prescribed by statistical theory, and in Sections 4.3 and 4.5 we give rules of thumb for their selection, following the recommendations of [52].

But it is also worth considering that in practice the hyperparameters may deviate from their optimal values, and here we study the sensitivity of the distortion estimates **H**^(*i*)^ on the radius *r* and rescaling *ϵ* hyperparameters, We repeat the analysis of Section 2.2.1, while varying the RMetric hyperparameters. We considered a grid of radius and rescaling *ϵ* parameters centered around the initial choice, *r* = 32.2 and *ϵ* = 5. We set *r* to 3 × the average neighborhood distance, as before. Our grid includes radii at 2^−1^, 2^−0.8^, …, 2^0.8^, 2 times the original value. We also tested 10 values of rescaling *ϵ* equally spaced in [3, 7]. For each combination of hyperparameters, we collected RMetric outputs **H**^(*i*)^ (*r, ϵ*) which govern ellipse size and orientation.

Fig 9 and Supplementary Figs 17, 18 give the results. Fig 9A - B show how the singular values of **H**^(*i*)^ (*r, ϵ*) vary across *ϵ* (x-axis) and *r* (color). These singular values govern ellipse size and eccentricity.

**Fig. 9:**
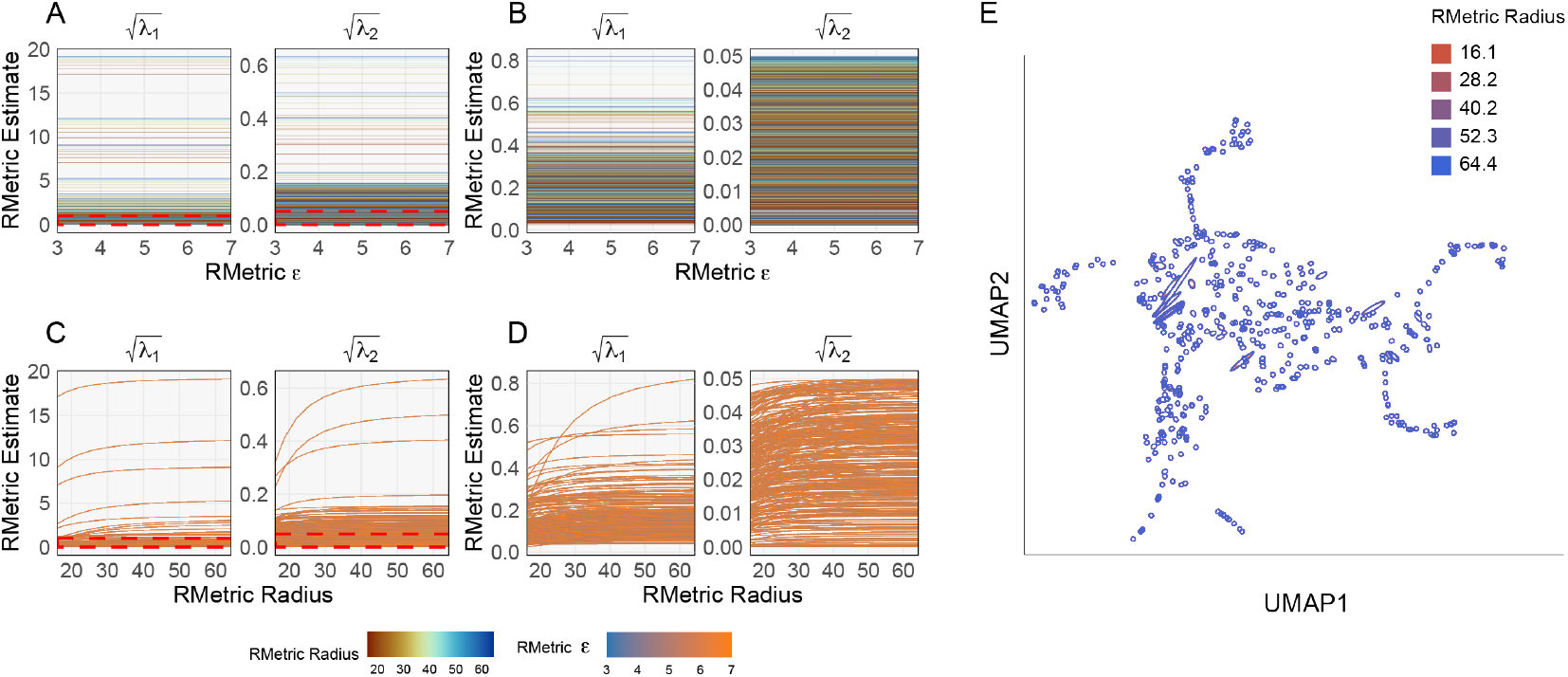
Dependence of ellipse visualization outputs on RMetric rescaling *ϵ* and radius hyperparameters. A. Singular values of **H**^(*i*)^ (*r, ϵ*) across *ϵ*. Each line corresponds to one sample and a fixed *r*. B. A version of panel A with *y*-axis ranges restricted to 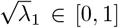 and 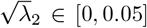, the region surrounded by the dashed red line. C. The analog of panel A with *r* along the *x*-axis and *ϵ* fixed. D. A version of panel C with *y*-axis truncated to the same range as (b). E. The RMetric estimated ellipses corresponding to random sample of 500 UMAP embedding points. Over each point, we draw multiple ellipses, corresponding to the radius and rescaling *ϵ* hyperparameters shown in the previous panels. Ellipse color encodes the radius parameter. See Supplementary Fig S18 for the analogous display with color encoding the rescaling *ϵ* hyperparameter.

We see that the singular values are extremely robust to the change in *ϵ*. Fig 9C - D provide the analogous views with radius on the *x*-axis and epsilon as color. Smaller radii lead to smaller ellipses, with larger ellipses showing the greater effect. Larger radii increase singular values until they plateau. This effect is expected, and it amounts to an error introduced by the small radius in the RMetricestimation; the details are beyond the scope of the paper. Supplementary Fig S17 gives the analogous visualizations for the singular vectors of **H**^(*i*)^ (*r, ϵ*), showing that most of these, too, are very stable across the RMetric hyperparameters. When singular values are nearly equal, they can switch ranks, causing their associated singular vectors to swap, and these are the abrupt changes in Supplementary Fig S17. Since these ellipses are nearly circular, this swap has little effect on the visualization.

Fig 9E shows a random sample of 500 points from the mammoth embedding in Fig 5. For each embedding points, we show the RMetric ellipses **H**^(*i*)^ (*r, ϵ*) across the *r, ϵ* hyperparameter grid from panels A - D. They confirm that ellipse visualizations can change slightly with *r*, leaving qualitative conclusions unaffected. Fig 18 gives the analogous visualization for varying *ϵ*. Consistent with Fig 9C-D, the effect on the resulting ellipses appears minimal.

The `distortions` package provides functions `expand_geoms` and `metric_sensitivity` for sensitivity analysis. The first function takes an RMetricgeometry object together with a grid of *r, ϵ* sensitivity factors, creating new RMetricgeometries associated with updated hyperparameters across the range specified by the input. The `metric_sensitivity` function computes the **H**^(*i*)^ and singular decompositions across the grid returned by the `expand_geoms`. These functions let users confirm that key findings remain robust across hyperparameter choices.

#### 2.5 Comparison with neMDBD and Sleepwalk

Existing approaches to distortion analysis differ conceptually. For example, Riemannian approaches [52] return tensor summaries at each embedding point without supporting interactivity; methods like [48] provide interactivity but only over scalar summaries. We compare three representative approaches – interactive [48], perturbation-based [34], and geometric (`distortions`). We evaluate methods on a variable density Swiss roll, which presents two embedding challenges. Capturing manifold structure as one continuous sheet is challenging – embedding map often introduce accidental twists and tears. The density variation also challenges embedding methods [44].

We sampled 1500 points from the following generative mechanism,

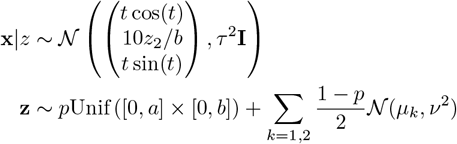

where *t* = 1.5*π*(*z*_1_*/a* + 1), *p* = 0.25, *a* = 2, *b* = 1, 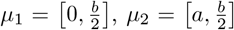, and *ν* = 0.2. The two Gaussian components concentrate density near the start and end of the roll. The uniform component ensures a baseline density throughout the roll. We sample versions of the data across a range of noise levels *τ* ∈ { 0, 0.5, 1, …, 3 }. We used *t*-SNE with perplexity set to 100 to embed each generated dataset into two dimensions. Fig 10A shows an embedding at *τ* = 0.5, revealing three types of distortion. High-density regions are spread across larger areas, a break appears near *t* ≈ 10, and a small twist occurs near *t* ≈ 8.

**Fig. 10:**
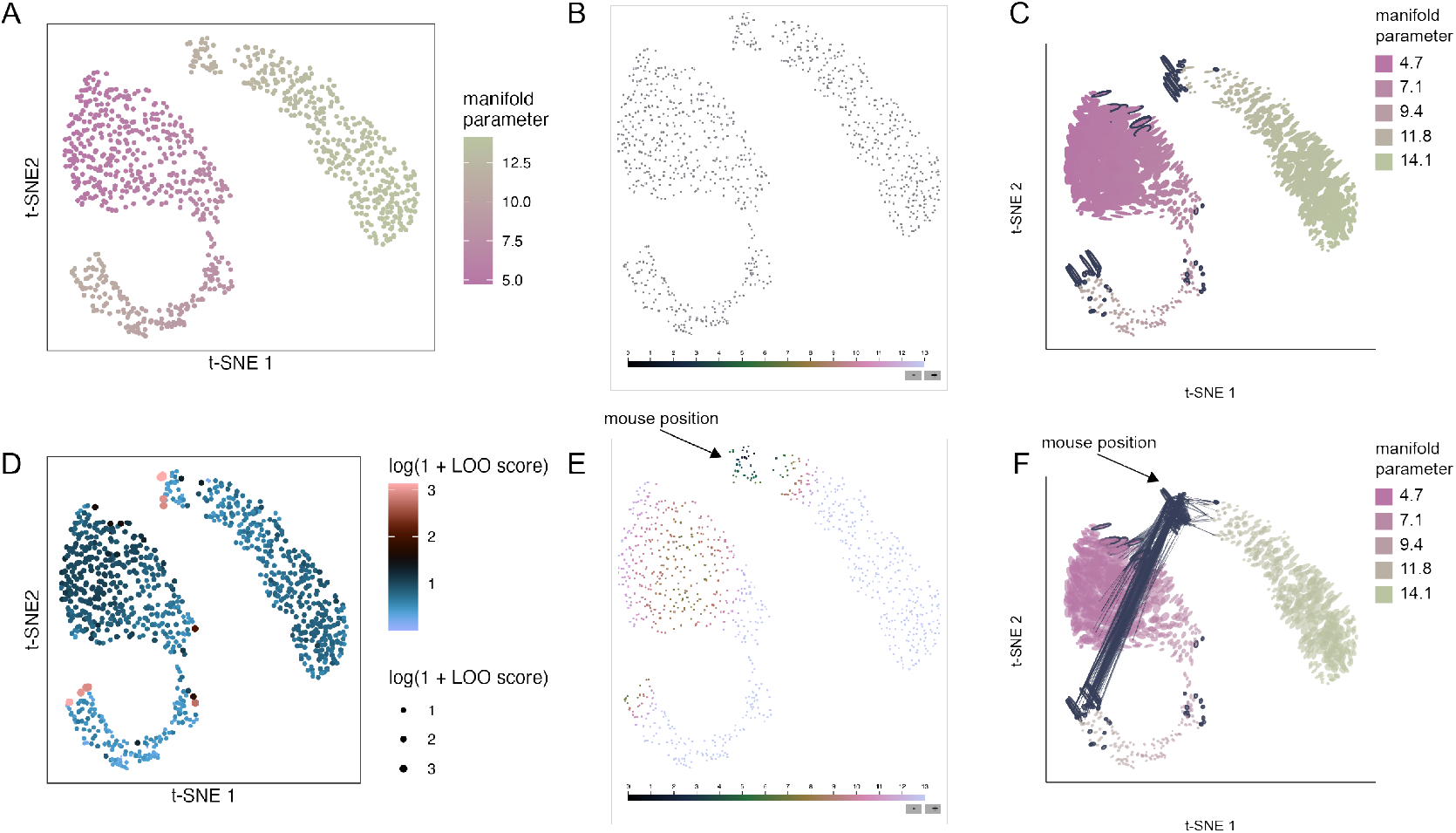
A. *t*-SNE embeddings of the variable density Swiss roll with *τ* = 0.5. The continuous roll has been broken into disconnected components, and high-density regions have been spread into wider lobes in the top-left and bottom-right of the scatterplot. B. Initial Sleepwalk view of the embedding. C. Initial `distortions` view of the embedding. D. LOO scores from the neMDDB algorithm when applied to the variable density Swiss roll with varying noise levels. E. Analogous results from the Sleepwalk algorithm. The cursor lies over a discontinuity. F. Analogous results for the fragmented neighborhoods visualization from `distortions`. Note the sensitivity to both changes in density (ellipse size) and fragmented neighborhoods (line segments).

We evaluate how two established baselines, neMDBD [34] and Sleepwalk [48], treat the embedding distortion. neMDB is based on a leave-one-out approach. Each point is embedded both jointly with all points and separately after freezing an embedding based on remaining points. Substantially different embeddings indicate high-distortion regions. The method output is a perturbation score describing the reliability of each point’s embedding. Sleepwalk, in contrast, is based on interactive visualization. After hovering over a point, the color of all points updates to show the distance to the hovered point in the original data space. Broken neighborhoods appear as points with low color distance that are far from the mouse cursor. We applied both methods with their defaults, except that in neMDBD we set `approx` to 2. This accelerates computation, potentially at a cost in optimization quality. This approximation minimally affects accuracy [34]. Even after this speedup, runtimes remain an order of magnitude higher than for Sleepwalk and `distortions` 2.

Fig 10 shows results at *τ* = 0.5. Supplementary Fig S19 - S23 show views for other *τ*. Fig 10D shows neMDBD’s LOO scores. Pink points reveal embedding discontinuities across the *t*-SNE components. Density changes appear more subtly through slightly larger LOO scores in the previously high-density regions. The twist apepars as dark-pink points on opposite sides in the middle of the gap of the left component. However, the true neighbors of the high LOO scoring points remain hidden, unlike in Sleepwalk or `distortions`. Sleepwalk highlights discontinuities and density changes in the embedding map (Fig 10B and E), but requires more viewer effort. The break appears when hovering near the right component’s edge. Points on the left boundary of the left component appear to be low distance to the cursor, though a large block of intermediate distance points with similar colors obscures them. The effect is also unclear in the static view, unlike neMDBD and `distortions`. Changes in density can be detected by attending to changes in the relative sizes of the points highlighted at a given distance. High-density regions that have been spread out are found by placing the mouse in regions that highlight larger embedding blocks in blue/green color. This demands more deliberate observation: comparing two views interactively taxes working memory more than the fixed encodings in neMDBD or `distortions`.

Fig 10C,F show the `distortions` view. Larger ellipses encode the spreading of high-density regions. In the static view, the fragmented neighborhoods are highlighted with dark-blue ellipse boundaries, and hovering these points reveals the particular fragmentation pattern. Encoding volume changes through size and fragmentation through overlaid links are more easily perceived than color gradients alone. When the noise level increases, the difference between the high leave-one-out scoring points and the bulk of insensitive points decreases. For example, consider *τ* = 2 − 3 in Fig 21, which have high LOO points more interspersed with lower LOO points, relative to the low *τ* case. This is not necessarily a problem – the embedding itself changes – but it makes views more difficult to parse. `distortions` also flags more points as distorted in these high-noise settings, but the interactive edges maintain interpretability. For example, the bottom-left and top right-corners of the embedding at *τ* = 2.5 are adjacent according to the Swiss roll parameter *t*, and the links connecting these regions when hovering over either corner makes this relationship clear. We have highlighted the corresponding Sleepwalk view in Supplementary Fig 20, which also gives evidence for the connection between the bottom-left and top-right corners of the embedding. However, this view required scanning the mouse over the entire view, rather than focusing in on regions with a large number of highlighted fragmented neighborhoods visible even in the static preview.

We note that Sleepwalk and RMetric are closely related through the inverse distortion (**H**^(*i*)^)^−1^. Indeed, since **H**^(*i*)^ is the distortion, its inverse is the respective *metric*. The ellipses corresponding to (**H**^(*i*)^)^−1^ will have approximately the same shape as the regions highlighted by Sleepwalk.

#### 2.6 Software architecture and extensibility

Considering that no single definition of distortion exists for nonlinear dimensionality reduction, the `distortions` package adopts a “loosely coupled” design to ensure extensibility [17]. Each visualization accepts viewer-provided specifications of fragmented neighborhoods or links. Alternative distortion metrics can be implemented in independent functions as long as their formats are consistent. Similarly, visualizations can composed from viewer-specified graphical marks and interactions, similar in spirit to ggplot2 [73] and altair [68]. For example, consider the interactions with fragmented neighborhoods of the PBMC data in Fig 5E - F. If the distorted neighborhoods are stored in a dictionary `N`, then the interactive plot can be created with and the result will appear in a jupyter notebook cell.

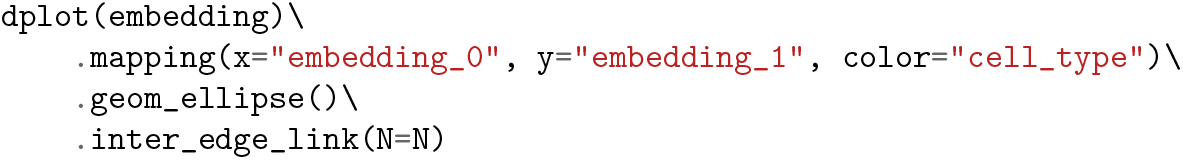

This loose coupling also simplifies the application of our visualization to other distortion summarization approaches. We illustrate this by using the scDEED algorithm [75] to flag dubious cells in a *t*-SNE embedding of the PBMC data (Fig 11A). This figure was generated by applying the default scDEED workflow to the PBMC data [1], adding local metrics **H**^(*i*)^ to the resulting embeddings, and replacing `embedding` and `N` in the call with the corresponding scDEED output. The resulting visualization is consistent with expectations – by hovering the mouse close to the scDEED flagged cells, we see the these cells often have neighbors in the original space that are placed far apart in the embedding space (Fig 11B). We note that no such interactive display has previously been available for output from the scDEED package.

**Fig. 11:**
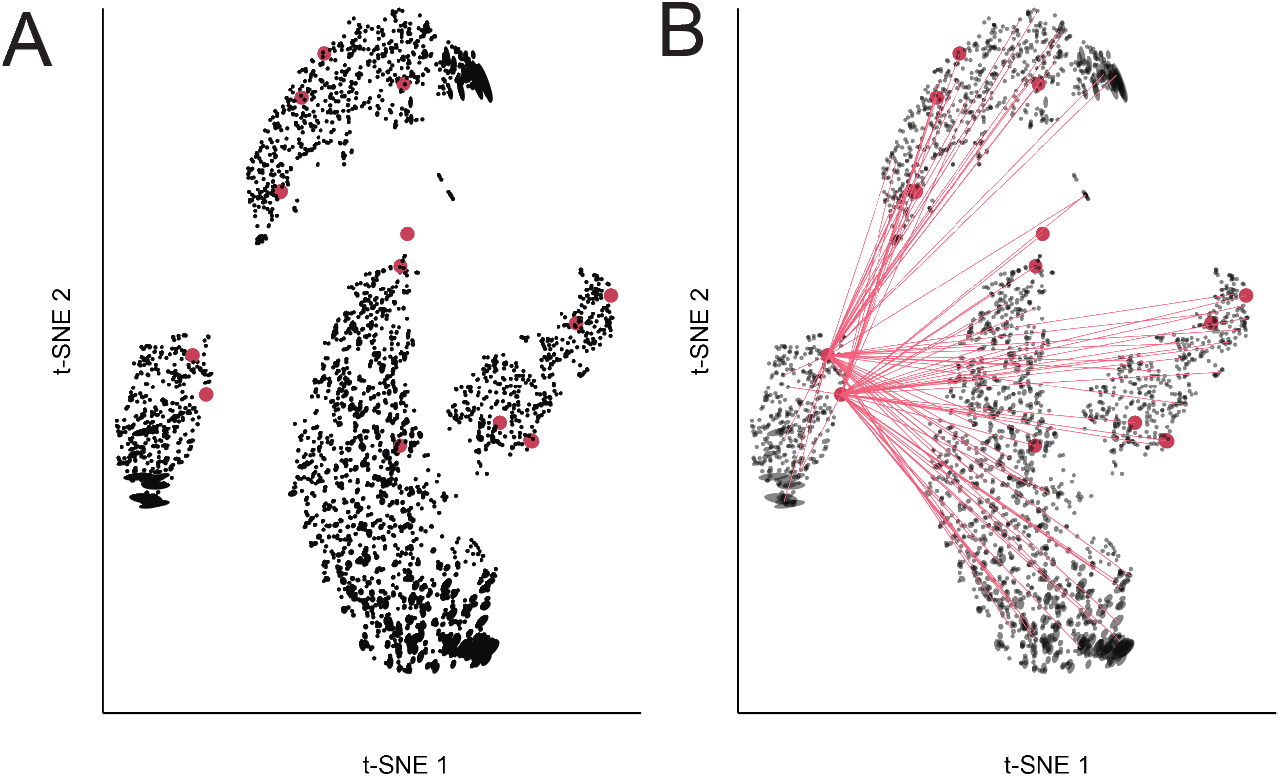
Integrating scDEED dubious embeddings into visualizations made with the `distortions` package. A. The PBMC data with dubious cells flagged by scDEED. B. Hovering over the far left cluster reveals that the scDEED flagged cells have neighbors lying across multiple cell types that are distant in the embedding space. Our visualization functions are designed to accommodate alternative definitions of nonlinear embedding distortion.

We can also customize the graphical marks, styling, and labels in a format familiar to to ggplot2 and altair users. For example, the visualization from the code block above can be customized using

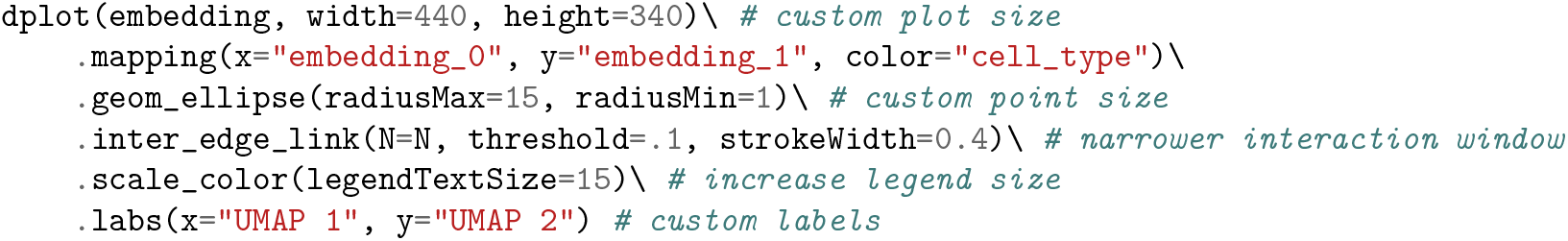

Further, we can switch from the fragmented neighborhood to the boxplot interaction (Fig 5D) by simply substituting the `inter_edge_link` call with `inter_boxplot`. This modular approach also enables the specification of new graphical marks and interactivity. For example, for large datasets, ellipses can be replaced with line segments, as in Fig 8D - E. This more compact encoding of local distortions is accomplished by substituting the `geom_ellipse` mark with `geom_hair`.

### 3 Summary and conclusions

Nonlinear embedding visualizations have been essential to progress in high-throughput biology, offering visual overviews that have guided advances in diverse applications like cell atlas construction [12, 54], cell differentiation trajectories [6, 16], and functional diversity mapping in metagenomes [61]. However, their potential for misinterpretation is well-documented [13, 72, 7, 26]. The community has made significant progress in characterizing and minimizing distortion [44, 75, 31, 34, 22], and the `distortions` package offers an interactive visualization toolbox that draws from manifold learning concepts and complements these advances.

Moderate distortions are accurately characterized by the RMetric algorithm, whose results can be graphically encoded in ellipse or hair plots, while more severe distortions are flagged via fragmented neighborhood plots. RMetric emphasizes how the intrinsic geometry is warped across different regions of the embedding space, alerting analysts to failures in density preservation and compression/dilation in certain embedding directions (Fig 2).

The fragmented neighborhoods function detects both embedding discontinuities (nearby points embedded far apart) and collapses (distant points embedded too close together). The first failure mode is more commonly observed and addressed in literature, but the second remains possible, as seen with the mammoth embedding (Fig 5A-C). Further, these issues extend to larger subsets of points, and we observed clusters that are more closely related in the original data space than the embedding suggests. In the PBMC fragmentation example (Fig 5D-F), we found that a subset of T cells had many neighbors coming from the monocyte cluster, despite these clusters appearing on the opposite sides of the embedding visualization. Further, interaction with distortion metrics highlighted trade-offs between the types of distortion introduced by different hyperparameter choices (Fig 6). Finally, local isometrization offers the scientist a kind of magnifying glass into the local geometry of the original data, making it possible to zoom in and query low-level sample relationships that can be lost in global reductions.

The summaries on fragmented neighborhoods depend on viewer-specified hyperparameters, like the number of bins *L* or neighborhood fraction *κ* in the bin-based definition. This can be seen as a limitation, and we mitigate it by setting reasonable default values. We recommend that the user takes advantage of these parameters under their control to explore more thoroughly neighborhood fragmentation, as these can seriously affect scientific interpretation of the embedding.

Regarding distortion, while the local distortion RMetric is a well-defined differential geometric quantity, measuring distortions at larger scales is open to subjective preferences. For example, the fragmented neighborhood definition relies on distances in the original high-dimensional space, while alternatives could consider geodesic distances along the original manifold embedded in the high-dimensional spaces. Pairs of points could be flagged by outlierness, or by constructing a neighborhood graph in the embedding space and comparing the two.

This package did not aim to exhaustively cover the existing embedding distortion measures. However, our modular software architecture will support straightforward extensions to new definitions and visualization layers. We expect that continued effort in this space will result in visualization techniques that can allow analysts to gain valuable insights from exploratory overviews while contextualizing their inherent limitations. While nonlinear dimensionality reduction methods cannot fully preserve all metric properties from the original data space, these exploratory views can guide more appropriate interpretation, allowing scientists to communicate results confidently and avoid the pitfalls of false discoveries due to algorithmic artefacts. By overlaying quantitative summaries of the distortion introduced by embedding algorithms, the `distortions` package aids researcher intuition and facilitates critical evaluation of the embedding visualizations that have become standard in modern biological analysis.

### 4 Methods

#### 4.1 Notation

In the following, matrices will be denoted in bold uppercase letters, e.g. ***A***, vectors in bold lowercase, e.g. ***v***, vector and matrix elements by additional subscripts, e.g. ***A***_*ii*_*′*, and other scalars by unbolded Latin and Greek letters. The index *i* will be reserved for denoting the *i*^*th*^ data point, and it will be used as a superscript on vectors and matrices associated with it. Thus, the original data is ***x***^(1)^, … ***x***^(*i*)^, … ***x***^(*n*)^ ∈ ℝ^*D*^, where *n* is the sample size and *D* is the dimension of the data. The embedded data points are denoted ***y***^(1)^, … … ***y***^(*n*)^ ∈ ℝ^*d*^, where *d* ≤ *D* is the *embedding dimension*. Here we formally define the variables that underlie the algorithms in the `distortions` package. For more background on the statistical and mathematical basis of embedding algorithms, the reader is referred to the [41] review.

#### 4.2 Neighborhood graph

Embedding algorithms such as UMAP [39], Isomap [66], *t*-SNE [35], DiffusionMaps [11], or LTSA [70] each output different embeddings ***y***^(1:*n*)^, but they all start from the same data representation, which is the *neighborhood graph*. Specifically, the first step in embedding data as well as in analyzing an embedding is to find neighbors of each data point ***x***^(*i*)^. This leads to the construction of the neighborhood graph as follows. Every data point ***x***^(*i*)^ represents a node in this graph, and two nodes are connected by an edge if their corresponding data points are neighbors. We use 𝒩_*i*_ to denote the neighbors of ***x***^(*i*)^ and *k*_*i*_ = |𝒩_*i*_ | be the number of neighbors of ***x***^(*i*)^. This graph, with suitable weights that summarize the local geometric and topological information in the data, is the typical input to a nonlinear dimension reduction algorithm.

There are two usual ways to define neighbors. In the *k-nearest neighbor (k-NN) graph*, 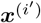 is the neighbor of ***x***^(*i*)^ iff 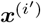 is among the closest *k* points to ***x***^(*i*)^. In a *radius-neighbor graph*, 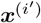 is a neighbor of ***x***^(*i*)^ iff 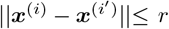, with *r* a parameter that defines the neighborhood scale. The *k*-NN graph has many computational advantages since it is connected for any *k >* 1 and each node has between *k* and 2*k* − 1 neighbors (including itself). Many software packages are available to construct (approximate) *k*-NN graphs fast for large data [20, 8, 43].

The distances between neighbors are stored in the distance matrix **A**, with **A**_*ii*_*′* being the distance 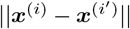 if 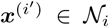 and infinity if 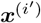 is not a neighbor of ***x***^(*i*)^. For biological data analysis, specialized distance functions can replace the generic Euclidean distance [32, 25, 77, 23]. From **A**, another data representation is calculated, in the form of an *n* × *n* matrix of weights that are decreasing with distances. This is called the *similarity matrix*. The weights are given by a *kernel function* [62], for example, the Gaussian kernel, defined as

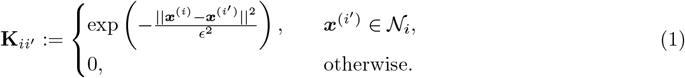

In the above, *ϵ*, the kernel width, is another hyperparameter that must be tuned. Note that, even if 𝒩_*i*_ would trivially contain all the data points, the similarity **K**_*ii*_*′* would be vanishingly small for faraway data points. Therefore, (1) effectively defines a radius-neighbor graph with *r* ∝ *ϵ*. Hence, a rule of thumb is to select *r* to be a small multiple of *ϵ* (e.g., *r* ≈ 3*ϵ*–10*ϵ*) [41]. ^1^

The neighborhood graph augmented with the distance matrix **A** or with similarity matrix **K** has many uses:

1. As stated above, it serves as a starting point for embedding algorithms.
2. In this paper, **K** is used to calculate the local distortion.
3. In this paper, **A** is used to detect the fragmented neighborhoods.
4. Neighborhood graphs are also used in estimating the intrinsic dimension, in Topological Data Analysis, namely in finding the loops and hollows in the data, as well as in other Geometric Data Analysis tasks.

While most embedding algorithms can take as input both types of neighborhood graphs (or resulting distance or similarity matrices), the embeddings obtained will be influenced by the type of graph and by the hyperparameter value used with it. For other uses, one type of graph or another may be optimal. In particular, for the purpose of estimating distortion, it is necessary to use the radius-neighbor graph, as this guarantees the distortion estimated is unbiased.

#### 4.3 Distortion estimation with the Dual Pushforward Riemannian Metric

The distortion estimation function `local_distortions()` implements the algorithm introduced by [51]. Given an embedding ***y***^(1:*n*)^ of data ***x***^(1:*n*)^ with similarity matrix **K** computed from radius-neighbor graph, `local_distortions()` outputs for each embedding point ***y***^(*i*)^ a *d × d* matrix **V**^(*i*)^ whose column 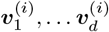 represent the *principal directions* of distortion at data point *i*. The stretch in direction 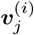 is given by 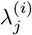. When 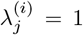 there is no stretch, for 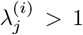 the embedding stretches the data in direction 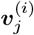, and for 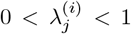 the embedding shrinks the data along this direction. Thus, the principal directions are orthogonal directions in the embedding where the algorithm induces pure stretch. Intuitively, the values 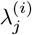 represents the local unit of length in direction 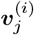.

The principal directions and stretch values result from the eigendecomposition of the symmetric, positive definite matrix 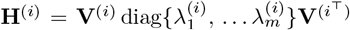 For an embedding with no distortion, namely an *isometric* embedding, **H**^(*i*)^ = **I**_*d*_ the unit matrix.

The local correction at ***y***^(*i*)^ is the inverse **G**^(*i*)^ of **H**^(*i*)^; in technical terms **G**^(*i*)^ is known as the *embedding (push-forward) Riemannian metric*. Obviously, the eigendecomposition of **G**^(*i*)^ is given by **V**^(*i*)^ and 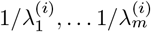. Thus, to correct the distortion in direction 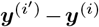, one calculates 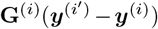. The orientation and length of this vector with origin in ***y***^(*i*)^ are the corrected direction and distance to nearby point ***y***^*′*^.

Hence, for any data embedding, it is sufficient to estimate, at all points ***y***^(1:*n*)^, the matrices **G**^(1:*n*)^, which represent the auxiliary information enabling correct distance computations, as if working with the original data, even though the embedding may not have preserved them. The same **G**^(1:*n*)^ can be used to preserve not only geodesic distances but also other geometric quantities such as angles between curves in ℳ or volumes of subsets of ℳ. Further uses of the distortion and correction matrices are described in [41, 51, 24], and here we present a corrected visualization based on **G**^(*i*)^.

#### 4.4 Computational complexity of RMetric

The complexity of the RMetric computation is dominated by the construction of the neighborhood graph. Since this graph is already computed for the purpose of embedding the data, we will only consider ^0^ the overhead. Obtaining the similarity **K** involves a fixed set of operations per graph edge (i.e. calculating the kernel value), hence order *m* operations total, where *m* is the number of edges in the neighborhood graph. Further computations also are proportional to *m*. Computing the RMetric at point *i* requires ∼ *k*_*i*_ *d*^2^ operations, where *k*_*i*_ is, as above, the number of neighbors of *i*. Hence, obtaining the RMetric at all points requires ∼ *md*^2^ operations.^2^ Further eigendecompositions and inversion of **H**^(*i*)^ are order *d*^3^ per data point, hence *nd*^3^ total.

Since the optimal neighborhood graph is a sparse graph (since it should only capture distance to nearby points and ignore the distances to far-away points), *m* is much smaller than the maximum value *n*(*n* − 1)*/*2. In practice, on large data sets, we have always found that computing the RMetric is much faster than computing the embedding itself. The same is true for the isometrization algorithm, in which the overhead after RMetric computation is to apply a simple transformation to every embedded point.

#### 4.5 Selecting the hyperparameter *ϵ*

We recommend [41] for a tutorial on the choice of parameters *k* and/or *ϵ* (with *r* being a small multiple of *ϵ*). An automatic method for choosing these parameters, reminiscent of cross-validation, was introduced by [51] and can be found in the `megaman` package https://mmp2.github.io/megaman/.

As a general rule of thumb, if a neighborhood graph results in a good embedding, then the neighborhood scale is the appropriate one for the RMetric as well. Hence, if the embedding is obtained via a radius-neighbor graph, then the same graph, or same **K** matrix should be used for `local_distortions()`. If a *k*-NN graph was used, then we recommend selecting *ϵ* so that the row sums of **K** average *k*, the neighborhood parameter of the *k*-NN graph.

#### 4.6 Identifying fragmented neighborhoods

To compare distances across the original and embedding space, let:

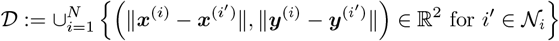

The `distortions` package supports two strategies for flagging neighbors with poorly preserved distances, which form the basis for defining fragmented neighborhoods.

##### Bin-based strategy

This approach partitions the original space distances into *L* evenly-sized bins and detects outliers in the embedding distances within each bin. Let *π*_*O*_ (𝒟) and *π*_*E*_ (𝒟) extract the original and embedding distances from 𝒟, respectively. With *d*_min_ = inf *π*_*O*_ (𝒟) and *d*_max_ = sup *π*_*O*_ (𝒟), set the binwidth 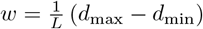 and partition the original data distances into intervals *I*_*l*_ = [*d* _min_ + *w* (*l* − 1), *d* _min_ + *wl*). The embedding distances within bin *l* are,

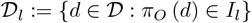

where we have abused notation and applied the projection *π*_*O*_ to an individual distance tuple *d*. For each bin, we compute the interquartile range of associated embedding distances,

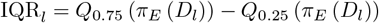

where *Q*_*α*_ extracts the *α*-quantile. A distance tuple *d* ∈ 𝒟 is considered outlying if,

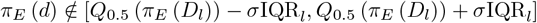

where *σ* is controls the outlier threshold. Note that neighborhood distances can be considered outlying for two qualitatively different reasons. The embedding distance may be either too large, where truly neighboring points may be artificially spread apart. This is labeled 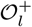 in Fig 12. Alternatively, they may be too small, where distant points are inappropriately collapsed on top of one another (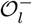 in Fig 12). All bin-*l* outliers are collected into the set 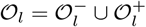.

**Fig. 12:**
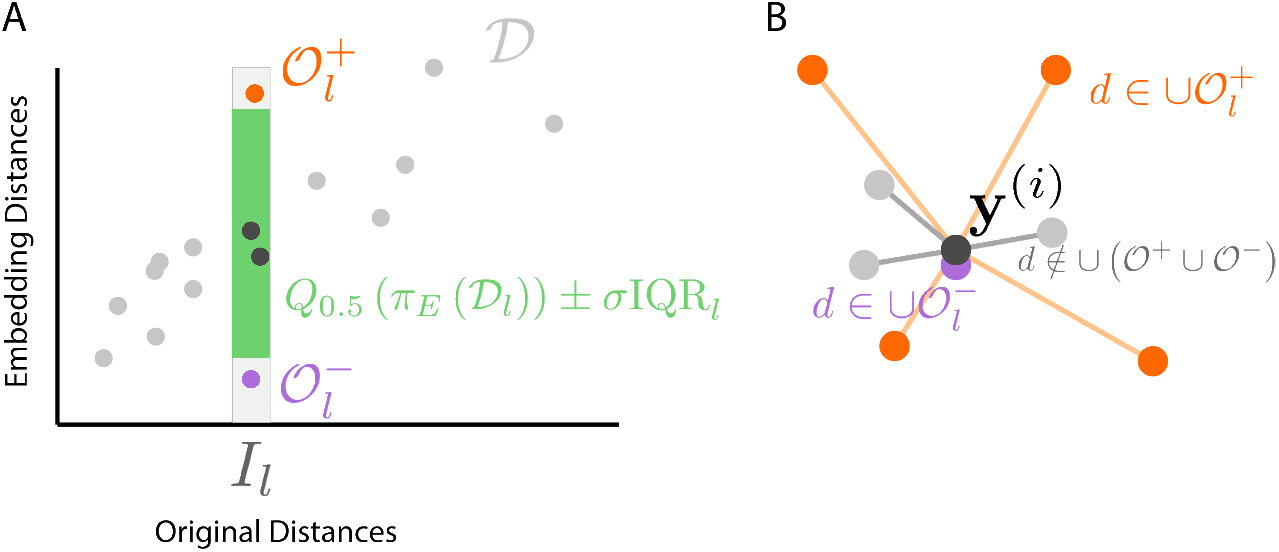
A graphical illustration of strategies used to flag fragmented neighborhoods. A. In the bin-based strategy, the original distances are partitioned into evenly-sized intervals *I*_*l*_. Within each bin, the interquartile range of embedding distances is computed. Original vs. embedding distance pairs that do not fall within a factor of *σ* times the IQR of the median embedding distance for that bin are flagged as outliers 𝒪_*l*_. B. In either the bin or window-based strategies, samples with many neighbor links belonging to ∪_*l*_ 𝒪_*l*_ are flagged as being the center of a fragmented neighborhood.

**Fig. 13:**
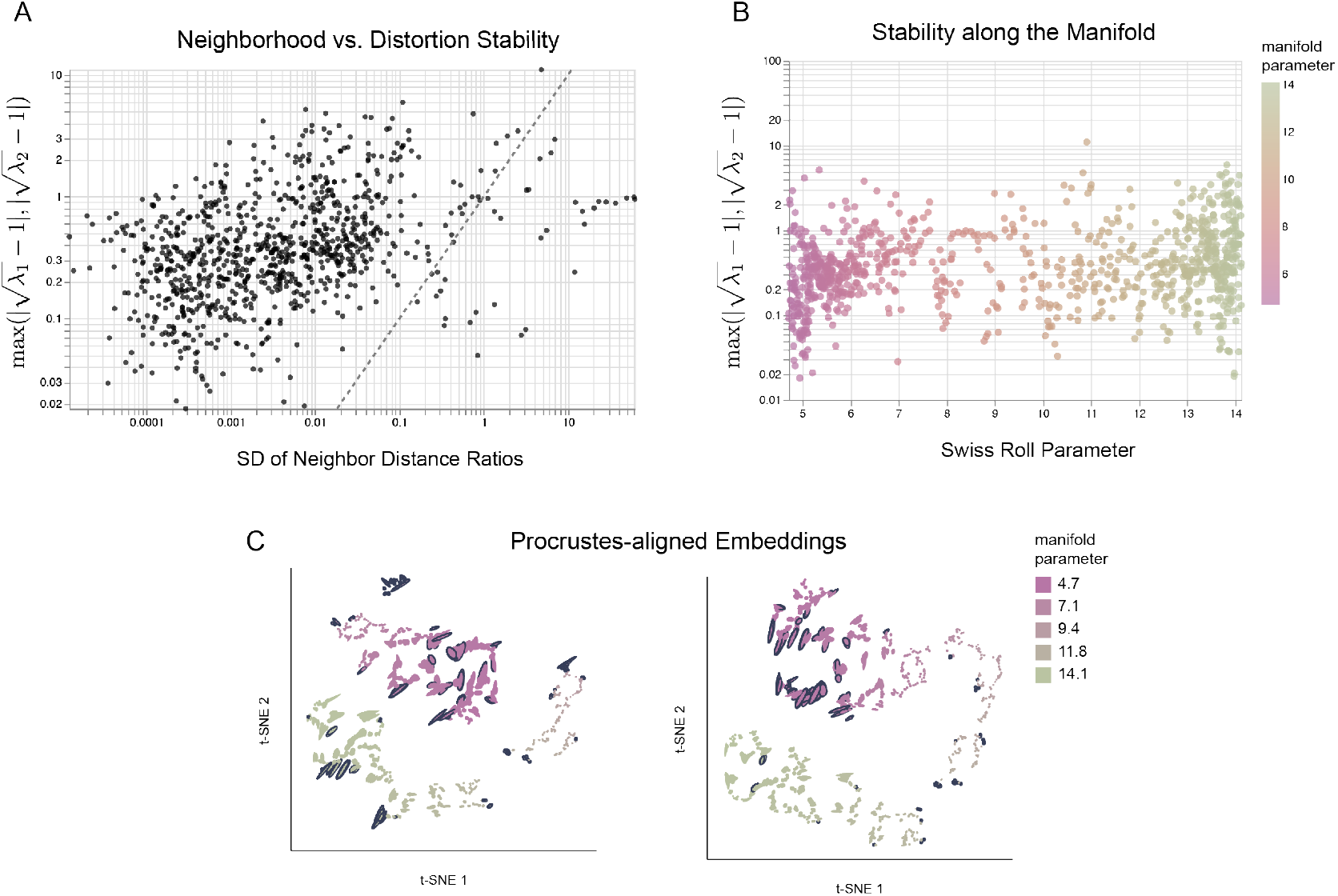
Stability across *t*-SNE initializations. A. The *x*-axis is embedding stability, measured by the variance in ratios of neighbor-neighbor distances across embeddings derived from two initializations. The *y*-axis quantifies RMetric instability using singular values of 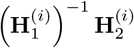. Embedding and RMetric stability are positively associated. B. RMetric instability across the Swiss roll parameter *t*. C. The *t*-SNE embeddings after Procrustes alignment.

We define fragmented neighborhoods using the outlier sets 𝒪_*l*_. We consider ***y***^(*i*)^ to be the center of a fragmented neighborhood if,

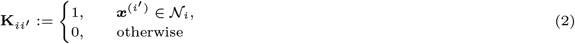

that is, if at least a fraction *κ* of the distances to its neighbors belong to at least one outlier set 𝒪_*l*_. This procedure is illustrated graphically in Fig 12.

##### Window-based strategy

The window-based strategy parallels the bin-based approach but uses running windows centered at each point. For each *d*_0_ ∈ 𝒟, we define a window 𝒟_win_ of the Δ nearest points with respect to *π*_*O*_ (*d*_0_). Within each window, we compute the interquartile range (IQR) of the embedding distances and flag *d* ∈ 𝒟 as an outlier if its embedding distance is more than *σ* IQRs from the median embedding distance in the window,

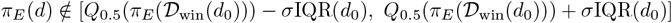

where IQR(*d*) is the interquartile range of *π*_*E*_(𝒟_win_(*d*)). As with the bin-based strategy, a neighborhood is fragmented if at least a fraction *κ* of its neighbor pairs are flagged as outliers. This approach leads to smoother IQR boundaries compared to the bin-based approach, but is more computationally involved.

#### 4.7 Focus-plus-context visual interaction

Adding distortion information to standard nonlinear embedding visualizations is challenging because the additional context can overwhelm an already complex visualization, making them even more difficult to understand. The `distortions` package addresses this challenge through the focus-plus-context principle [18, 28, 55]. This approach displays distortion information locally (“focus”) while maintaining the broader visual overview (“context”). The region within which to display additional information is set by the viewer’s interactions. We implement three forms of focus-plus-context interactivity, adapted to visualize fragmented neighborhoods, distance preservation, and local isometries, respectively.

##### 4.7.1 Mouseover interactions to reveal fragmented neighborhoods

This visualization supplements the original embedding overview by highlighting fragmented neighborhoods when their centers are hovered over. The centers may be defined using either the bin-based or window-based strategies described above. Before interaction, the fragmented neighborhood centers are highlighted with a distinctive stroke and color, guiding attention to regions of the embedding that are enriched with fragmentation. When the viewer’s mouse is moved to a location *m* ∈ ℝ^2^, all fragmented neighborhoods with centers within a distance *δ* of *m* are highlighted. Specifically, an edge is drawn between ***y*** ∈ ℝ^2^ and ***y***^*′*^ ∈ ℝ^2^ if:

1. ∥ ***y*** − *m* ∥ ≤ *δ*.
2. ***y*** satisfies the fragmented neighborhood criterion (Equation 3).
3. ***y***^*′*^ is one of the top *k* neighbors of ***y*** in the original data space.

The neighbors ***y***^*′*^ are highlighted when their corresponding edge links are visible. The hyperparameters *δ* and *k* must be specified by the viewer. We default to the *k* used in the original embedding. Since the neighborhoods are fragmented, the associated edge links typically span large regions of the embedding space, making interactive updates necessary to prevent occlusion from overlapping edges.

##### 4.7.2 Brush interactions to visualize distance preservation

The focus-plus-context principle supports visualization of individual edges with poorly preserved distances, rather than entire neighborhoods. A brushable widget is placed alongside the main embedding visualization and displays boxplots that compare binned distances in the original data space (*x*-axis) with the distances in the embedding space (*y*-axis). This boxplot overview builds on the static approach of [26]. Boxplot whiskers are capped at *σ* times the IQR, with outliers beyond this range drawn as distinct points. The number of bins and *σ* are user-specified hyperparameters. As the brush is moved, the embedding visualization updates to highlight edges between neighbors with brushed and outlying embedding distances. The coordinated display allows viewers to focus on specific distorted neighbor pairs within the context given by the overview boxplots.

##### 4.7.3 Mouseover interactions to update local isometries

The `distortions` package supports interactions that provide an intuitive understanding of local metric differences induced by the embedding. In this view, the mouse’s position is used to isometrize neighborhoods centered around it, providing an interactive, local version of the isometrization algorithm from [51]. Rather than modifying the entire embedding to induce an isometry around a selected point, this view updates the region around the mouse position. To isometrize the embedding with respect to sample *i*^*′*^, [51] suggest the transformation,
d

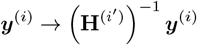

For focus-plus-context interaction, we isometrize only samples near the mouse position *m* and smoothly interpolate the transformation as the mouse moves between samples. We implement,

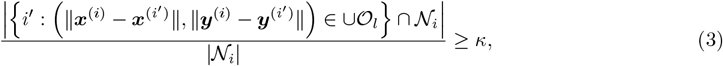

where,

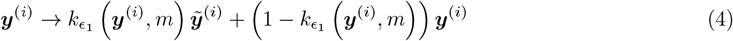

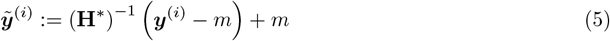

and 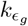 denotes the Gaussian kernel with bandwidth *ϵ*_*g*_ and **H**^∗^ represents a local average of **H**^(*i*)^. The parameter *ϵ*_1_ controls the size of the region affected by isometrization, and *ϵ*_2_ controls the region defining **H**^∗^. This interactive coordinate system update is related to fisheye distortion [56], where local geometries are deliberately distorted to focus on specific samples.

#### 4.8 Exporting corrected embeddings and static views

In the local isometrization visualization, the coordinates, sizes, and orientations of sample ellipses change in response to mouse interaction. In addition to interactive exploration of these corrected embeddings, it is also of interest to use these embeddings in downstream tasks, like cell clustering or marker gene analysis. To support these downstream applications, the `distortions` package provides methods for exporting the coordinates from any fixed view into a pandas DataFrame, which can also be saved as CSV. To freeze the current view, the viewer can double click or press the Escape key. While the view is frozen, any mouse interactions have no effect on the embedding. Calling `.correct` on the associated static view will return a pandas DataFrame with coordinates, orientations, and sizes of the underlying ellipses.

In addition to corrected embedding coordinates, the package allows viewers to export static views in scalable vector graphics format (SVG). This is done by freezing the view and then calling the `.save` method with the output file path as the argument. Multiple views can be saved in sequence by repeated unfreezing, interaction, freezing, and saving. The associated SVG files can be edited in standard vector graphics editors like Adobe Illustrator or Inkscape. Examples of the appropriate use of `.correct` and `.save` can be found in the “Exporting Results” article in the online documentation.

#### 4.9 Package software architecture

The `distortions` software architecture must support low-level graphical marks, like ellipses, and interactions, like updating fragmented neighborhood links on mouseover, that are unavailable in existing visualization software. These customizations cannot come at the cost of support for higher-level data structures from modern computational biology software. To this end, we have defined a standalone javascript package (`distortions-js`, https://www.npmjs.com/package/distortions) for visual components and interactions, and a separate python package (`distortions`, https://pypi.org/project/distortions/) for higher-level algorithms and distortion computation. The javascript implementation is built around a `DistortionPlot` class, which exports a `mapping` method to encode dataset fields in the visual channels from each `geom*` element, as well as methods for each interaction type. All graphical marks are rendered as SVG elements on a parent canvas. This is necessary, as standard javascript plotting libraries like `vega` [58] and `observable plot` [50] do not support ellipse visualization. Brush events are implemented using the d3-brush library [4], and legends are drawn using d3-legend [33].

The python package connects to `distortions-js` through the `anywidget` [36] package, allowing interactive javascript execution within Jupyter and Quarto notebooks. This approach converts python dictionary objects storing data and plot specifications into javascript data structures for visualization in the browser or notebook cell. The embeddings can be passed in through an `AnnData` experimental object [74]. For intrinsic geometry estimation, we use the `megaman` package [40], which is designed for scalable nonlinear dimensionality reduction and supports estimation of the local metrics **H**^(*i*)^ for each sample *i*. Open source code can be found at https://github.com/krisrs1128/distortions and https://github.com/krisrs1128/distortions-js, documentation is given at https://krisrs1128.github.io/distortions/site/. We note that the packages can be used independently.

#### 4.10 Key Points

- The `distortions` package applies the RMetric algorithm to biological data for the first time, showing how embedding algorithms warp neighborhood distances.
- Embedding methods necessarily distort distances, but local isometrization can interactively correct warping effects within a region-of-interest.
- Neighborhood fragmentation and anisotropy visualizations help to distinguish biological signal from technical embedding artifacts in both single-cell atlas construction and differentiation analyses.
- Distortion visualizations reveal the magnitude and nature of distortions, clarifying trade-offs between algorithms and hyperparameters.

## Data Availability

All data used in this study are publicly available and analysis can be reproduced using the code on the package documentation homepage https://krisrs1128.github.io/distortions/site/.

## A Appendix

### A.1 Working with large-scale data

Single-cell studies now routinely profile millions of cells. Visualizing the associated nonlinear embeddings using scatterplots can use to severe overplotting, and the ellipse marks used by `distortions` potentially exacerbate this issue, since viewers are expected to compare the orientations and sizes of ellipses, rather than just cell-wise coordinates. For work with the `distortions` package, we recommend learning both the embeddings and the local metrics **H**^(*i*)^ using the full data, since both steps only require nearest neighbors graphs, which can be computed or approximated efficiently. For visualization, it is challenging to perceive differences in overplotted data, even with high-resolution screens.

Therefore, we recommend randomly sampling up to this many points in a final embedding plot. The associated ellipses still reflect estimates of the local metrics made using the full dataset. Alternatively, if interest lies in a particular trajectory or subset of cell types, it may be possible to narrow the visualization to these more narrowly-defined subsets, as done in Section 2.3.3. The problem of identifying scalable alternatives to scatterplots has been extensively studied in the visualization literature. Unfortunately, since ellipse encodings are critical for communicating distortion in the `distortions` package, these scalable alternatives are not immediately applicable. Nonetheless, we expect that future research will identify improved data approximations or graphical encodings. For example, the original data can be replaced with randomized sketches [19, 67], and scatterplots can be replaced with density-based alternatives [2, 38, 9].

### A.2 Comparing embedding and neighborhood stability

Section 2.4 shows that RMetric varies smoothly across hyperparameters. We next study stability with respect to changes in the embedding. We applied *t*-SNE to the variable density, *τ* = 0, Swiss roll data (Section 10) using two random initializations and perplexity 10 to induce instability, aligning the results using a Procrustes rotation. Supplementary Fig S13C shows the embeddings with RMetricestimates overlaid.

For sample *i*, let 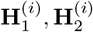 denote the RMetric estimates across the random seeds, and consider the eigenvalues of 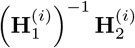, which we write as 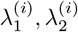. We use 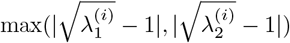 to measure the variability of **H**^(*i*)^ across seeds. This metric captures changes in ellipse size and eccentricity but is rotation invariant. Hence, it lower bounds the instability of RMetricat sample *i*. To quantify embedding stability, we formed ratios 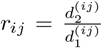 and 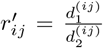 for all pairs of *K* = 15 nearest neighbors *i, j* in the original space. For each *i*, we compute the variances of *r*_*ij*_ and 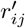 across neighbors *j*, taking the maximum as our stability measure. Supplementary Fig S13A relates these metrics. Most points lie above the identity, suggesting that RMetric is less stable than the embeddings themselves. The positive association indicates that RMetric instability tracks embedding instability. Supplementary Fig S13B highlights an association between RMetric and position along the roll, with more stability at lower ranges of *t*.

### A.3 Additional baseline comparisons

We compared `distortions` to neMDBD [31] and Sleepwalk [48] algorithms applied to the Gaussian mixture and PBMC data from Sections 2.1.1 and 2.1.2. To accelerate neMDBD on the PBMC data, we subsampled down to 1000 cells. We ran Sleepwalk and `distortions` on the full datasets. The corresponding runtimes are presented in Table 1.

**Table 1:** Runtime for methods considered in Supplementary Section A.3. As in Table 2, the neMDBD package was run with parameter approx=2.

| Method | Dataset | Seconds |
| --- | --- | --- |
| distortions | Gaussian Mixture | 0.181 |
| neMDBD | Gaussian Mixture | 1234.304 |
| sleepwalk | Gaussian Mixture | 0.016 |
| distortions | PBMC | 0.464 |
| neMDBD | PBMC | 1030.709 |
| sleepwalk | PBMC | 0.030 |

**Table 2:** Runtime for methods considered in Section 2.5, averaged across noise levels *τ*. The neMDBD package was run with parameter approx=2 to accelerate leave-one-out computation.

| Method | Mean (seconds) | SD (seconds) |
| --- | --- | --- |
| distortions | 0.93 | 0.05 |
| neMDBD | 1156.96 | 27.55 |
| sleepwalk | 0.04 | 0.02 |

In the Gaussian mixture example, mousing over the two mixture components reveals the density preservation failure. In the embedding space, the clusters appear similar in width, but hovering over the right-hand cluster highlights its smaller interpoint distances (Fig 15A-B). This is the small, high-density cluster from the original data. When using neMDBD, this high-density cluster is associated with larger perturbation scores (Fig 14). The high-density cluster has spread farther in the embedding space.

**Fig. 14:**
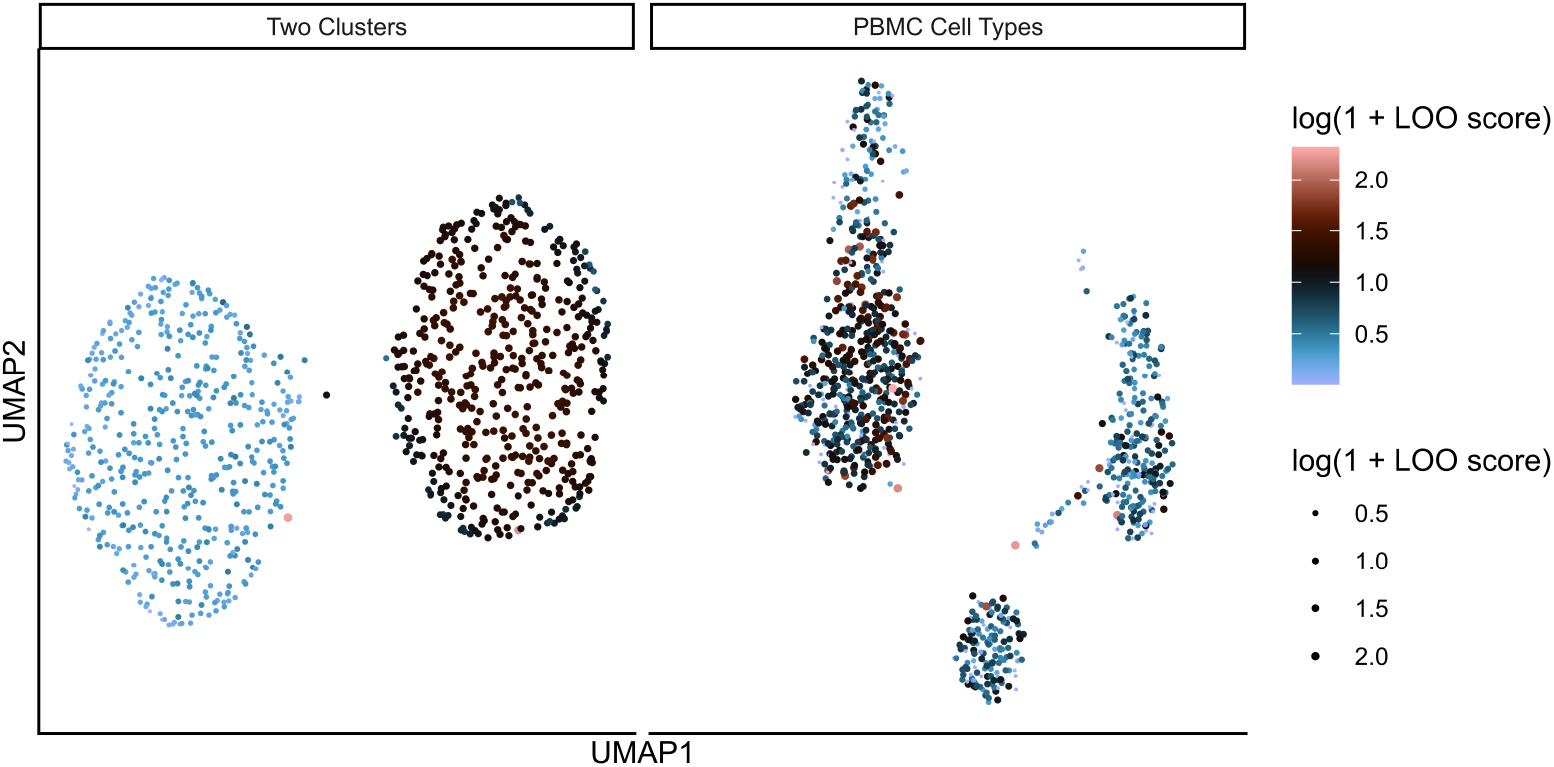
Perturbation scores from the neMDBD algorithm applied to the gaussian mixture simulation and the PBMC data from Sections 2.1.1 and 2.1.2, respectively. In the simulation example, the high density cluster has systematically higher perturbation scores. In the PBMC atlas, a subset of monocytes appears to have higher perturbation on average, and outlying perturbation scores appear near the cell type boundaries.

**Fig. 15:**
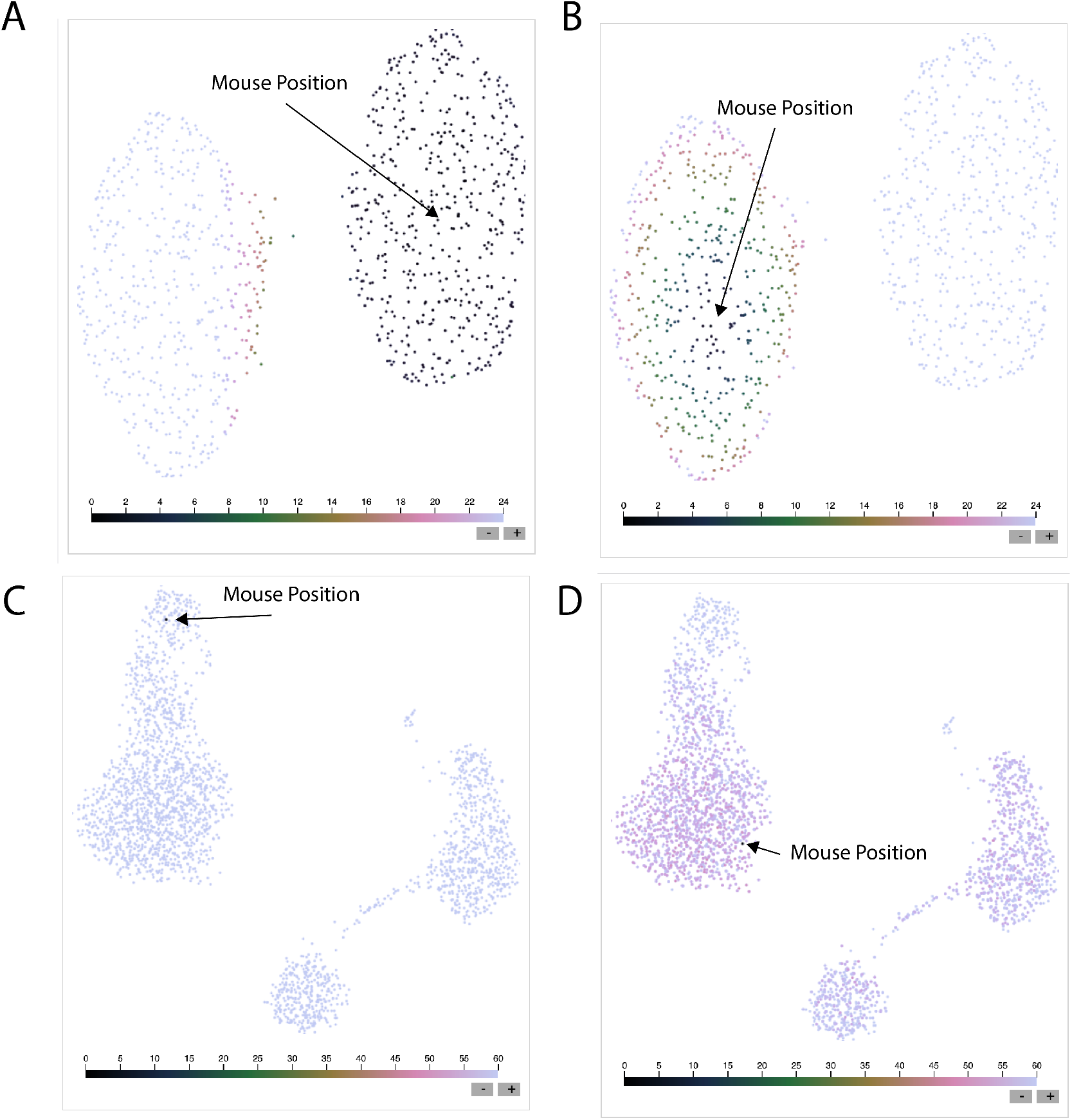
Sleepwalk applied to the Gaussian mixture and PBMC atlas examples. Between panels A and B, the mouse position switches from the high to the low density cluster. Between panels C and D, the mouse position switches from the dendritic to a subset of monocyte cell types.

### A.4 Supplementary tables and figures

**Fig. 16:**
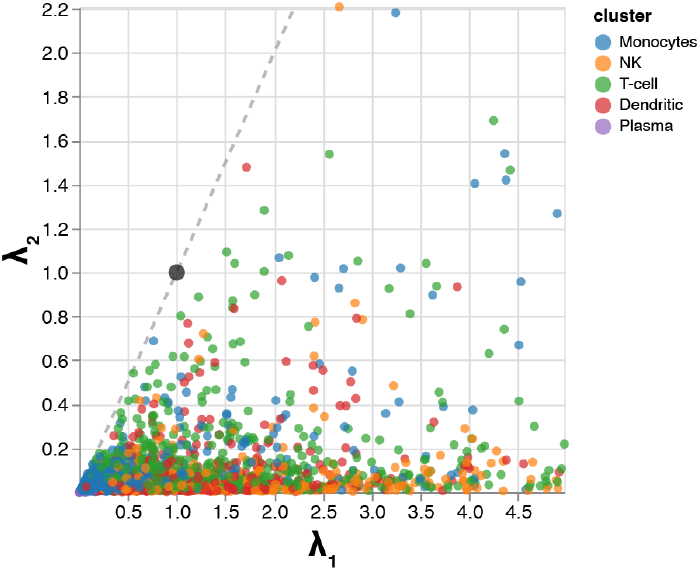
A zoomed-in version of Fig 4D. We have restricted to cells with 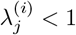. A second mode of smaller, less eccentric monocytes is visible in this view and contrasts with those that occupy the top right region of Fig 4D. We also see a small cluster of dendritic cells with singular values near the origin, corresponding to the small cluster placed near T cells in Figure 4A-C.

**Fig. 17:**
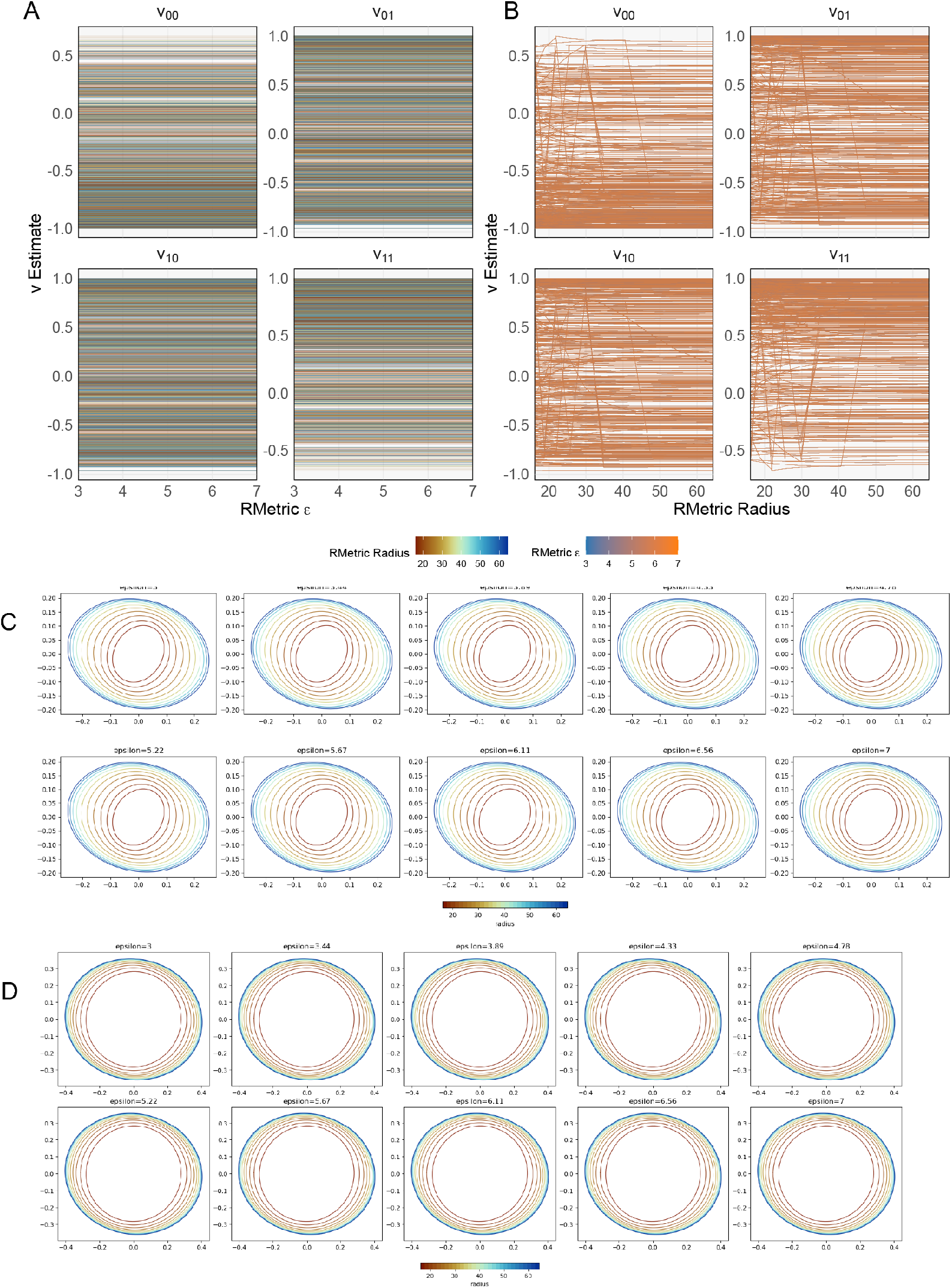
Sensitivity of singular vectors of **H**^(*i*)^ across hyperparameter choices. These are derived from the same experiment as Fig 9. Each line in panels A – B corresponds to one embedding point. While singular vectors appear insensitive to changes in *ϵ*, they sometimes appear to jump across small changes in radius. Panels C - D investigate these points more closely. These panels show the two samples with the largest absolute difference in singular vector coordinate values across neighboring values of *r*. Each subpanel within panel D shows the ellipse for one choice of radius and all values of *ϵ*. We note that both examples are ellipses whose eccentricitiy is near zero and where the ranking of the two singular vectors switches as the radius changes. We conclude that while the coordinates of the singular vectors may occassionally be unstable, the ellipses themselves change smoothly.

**Fig. 18:**
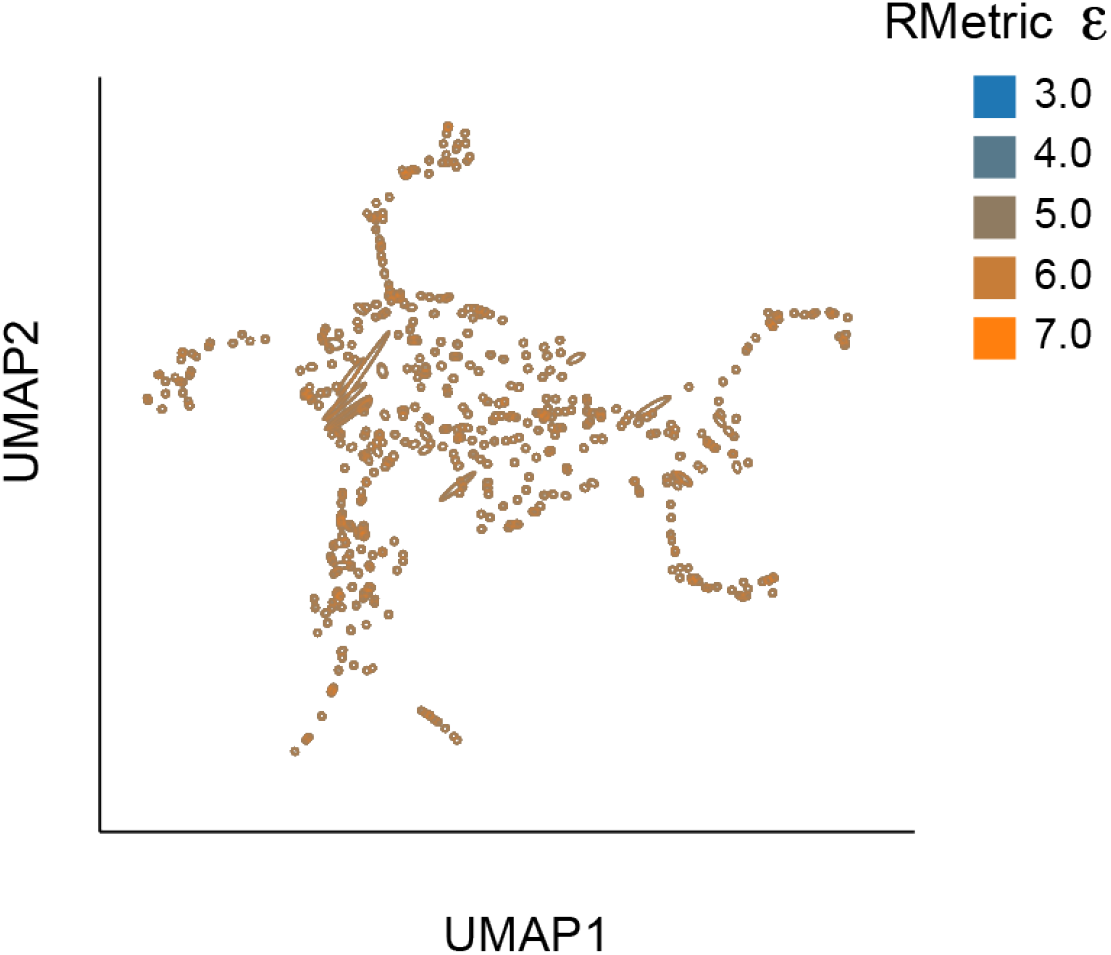
The analog of Fig 9E where ellipse borders encode the RMetric radius hyperparameter. Radius has a weaker effect on ellipse size and eccentricity than *ϵ*.

**Fig. 19:**
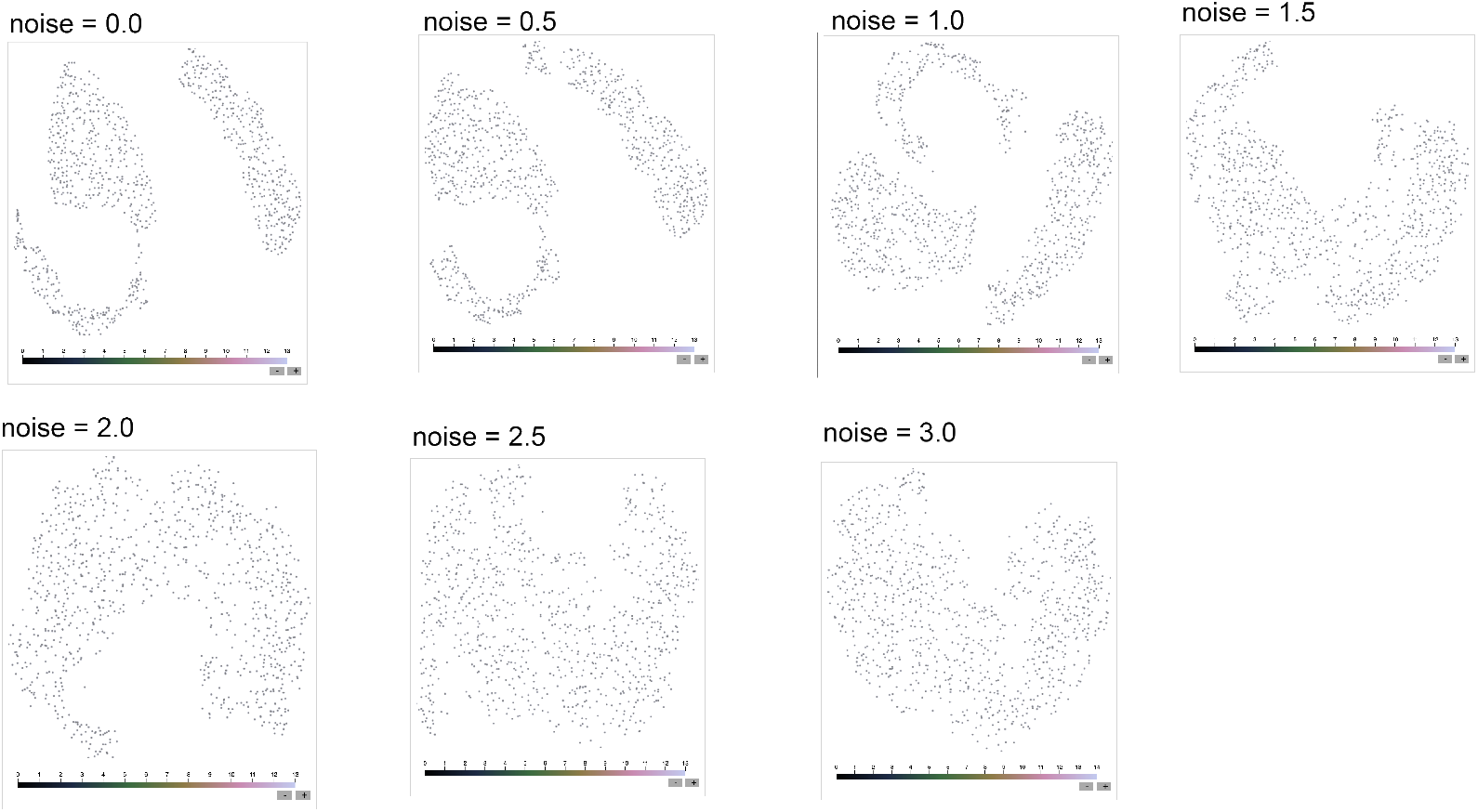
Sleepwalk visualization of the noisy, variable density Swiss roll across noise levels *τ*. Updated views after interaction are shown in Supplemental Fig S20.

**Fig. 20:**
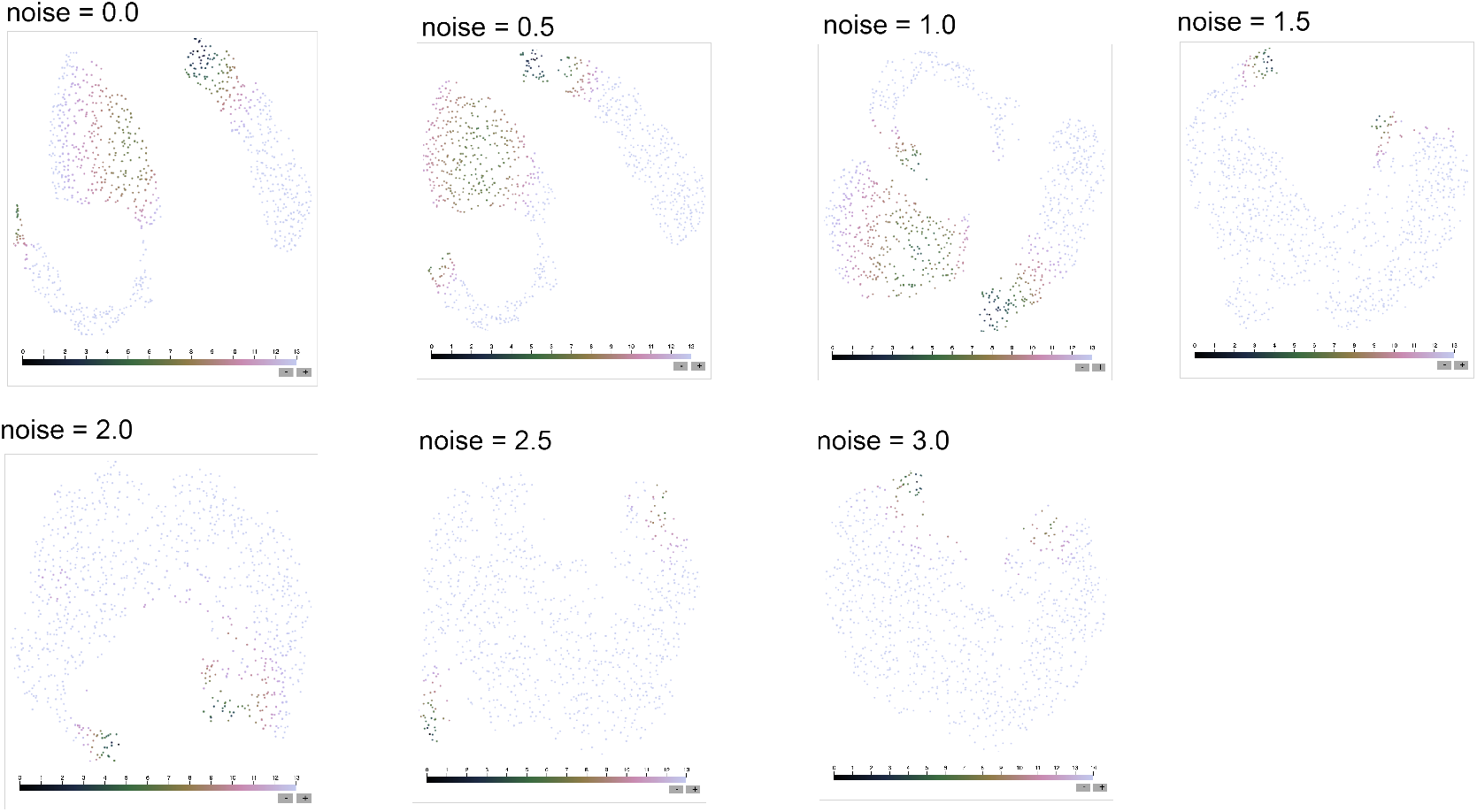
A version of Supplemental Fig S19 after placing the Sleepwalk cursor over a fragmented region of the *t*-SNE embedding. Darker blue points are close the nearest hovered point in the original data space. The size of the embedding region with low distance to hovered point in the original data space reflects a failure to preserve density.

**Fig. 21:**
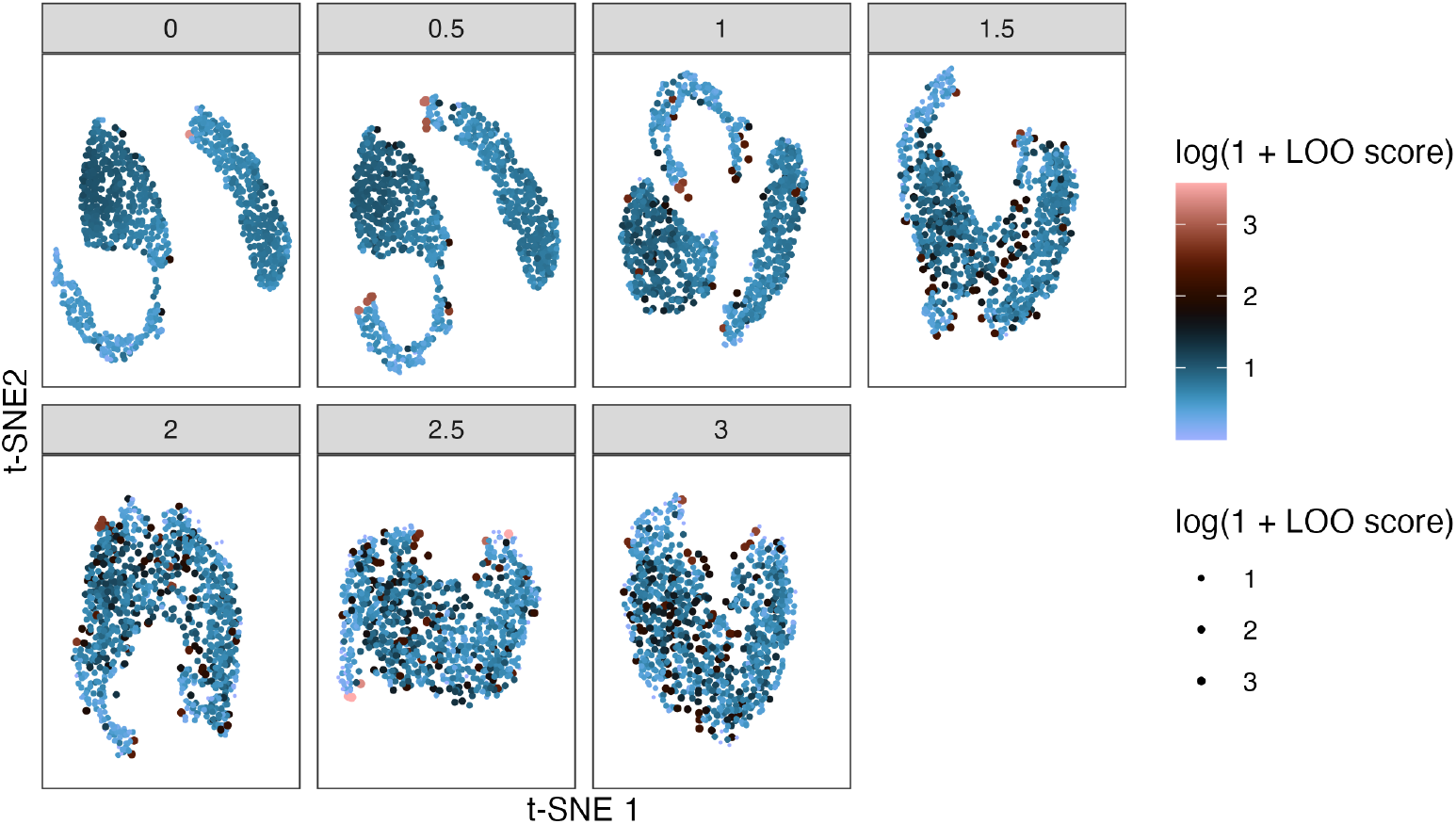
LOO perturbation scores applied to the noisy, variable density Swiss roll noise levels. Scores are shown on a log (1 + *x*) scale so that extreme outlier scores do not drown out other variation in LOO perturbation score. Perturbation scores slightly increase among points in the original data space’s highdensity regions. As the noise level *τ* increases, a larger fraction of points have above average perturbation scores, but the maximum scores are less extreme.

**Fig. 22:**
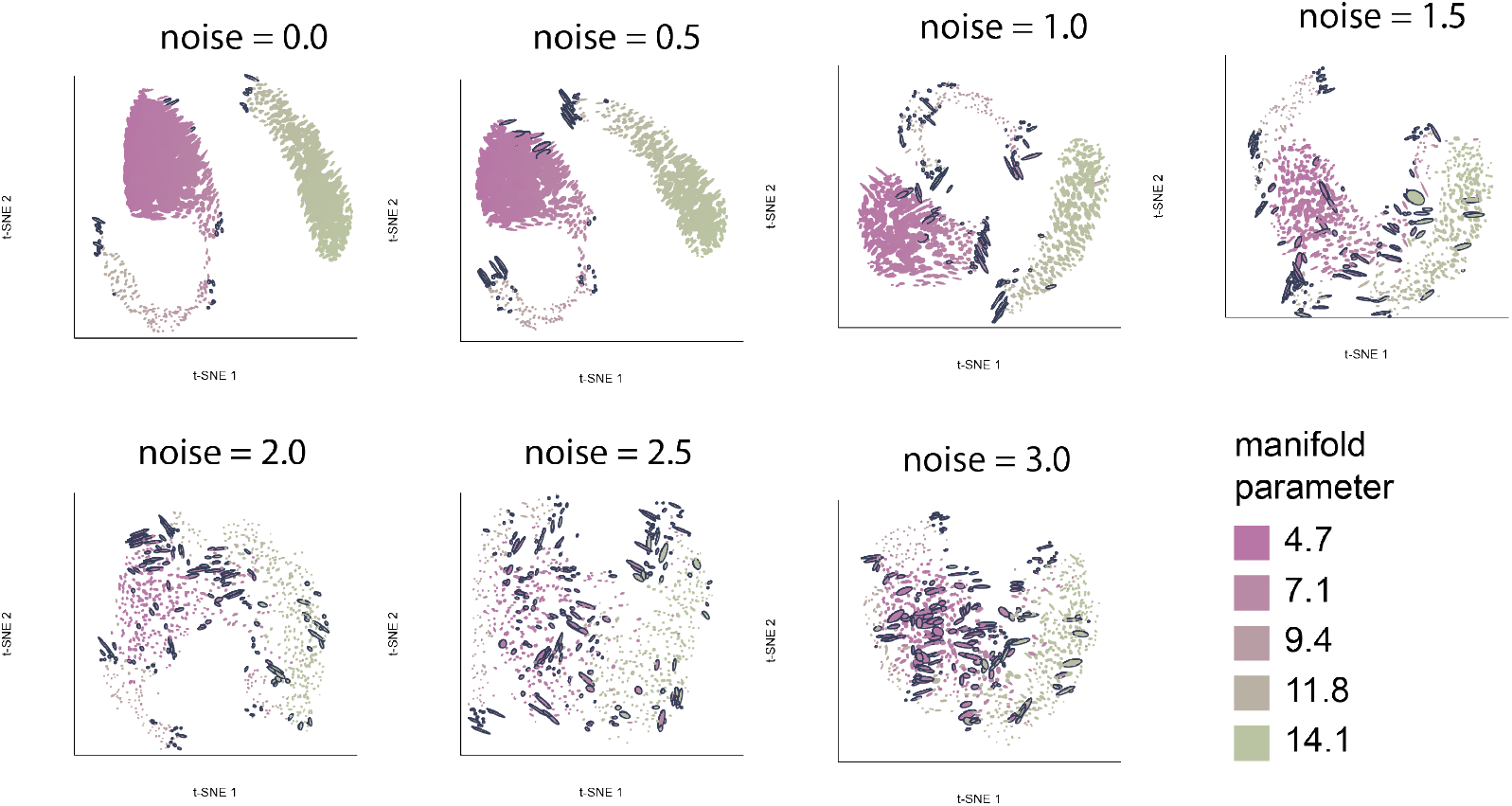
Initial views of the `distortions` fragmented neighborhoods visualization for *t*-SNE embeddings of the noisy, variable density Swiss roll, before placing the cursor over the panels. Difference in ellipse size reflect the failure for *t*-SNE to preserve the original data density in the embedding.

**Fig. 23:**
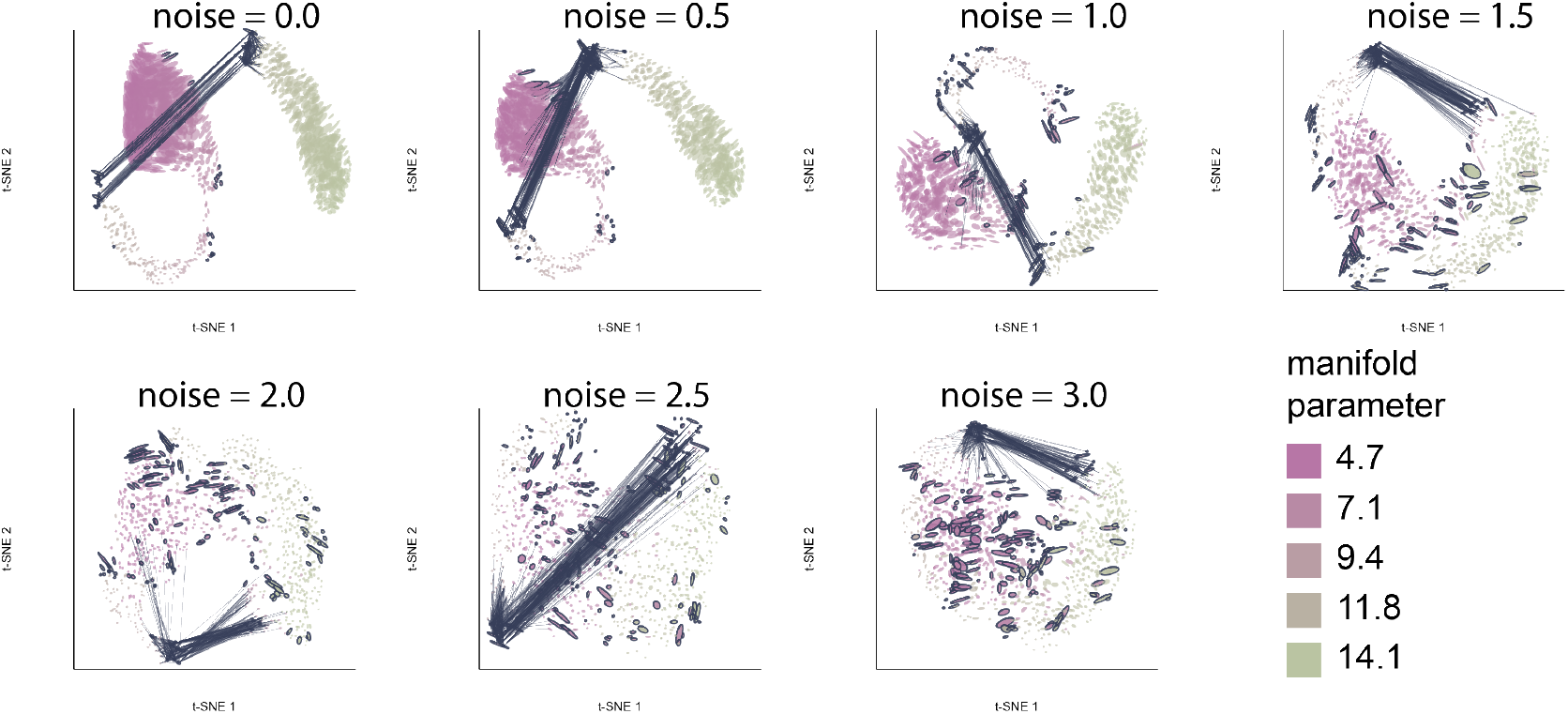
A version of Supplemental Fig S22 after hovering over example fragmented neighborhoods highlighted in the initial view. In addition to the density preservation failure evident in that view, we can identify discontinuities in the embedding map.

In the PBMC example, Sleepwalk’s color encoding occasionally spaces color so widely that all but the nearest neighbor point appear identical (Fig 15C). Hovering over other points appropriately highlights neighboring cell types (Fig 15D). The neMDBD algorithm identifies monocytes with elevated perturbation scores (Fig 14), matching the larger ellipses in Fig 4A from the `distortions` package. Boundary points between some cell types (bright pink) also draw attention to perturbation sensitivity.

1 Sometimes, the simple similarity 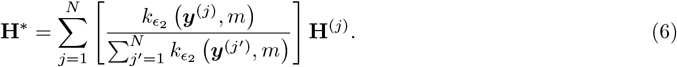 is used. This similarity matrix **K** is the unweighted adjacency matrix of the neighborhood graph, and completely ignores the distances.

2 Since ∑_i_*k*_*i*_ = 2*m*.

